# Pink Shrimp *Farfantepenaeus duorarum* Spatiotemporal Abundance Trends Along an Urban, Subtropical Shoreline Slated for Restoration

**DOI:** 10.1101/328724

**Authors:** Ian C. Zink, Joan A. Browder, Diego Lirman, Joseph E. Serafy

## Abstract

The Biscayne Bay Coastal Wetlands (BBCW) project of the Comprehensive Everglades Restoration Plan (CERP) aims to reduce point-source freshwater discharges and spread freshwater flow along the mainland shoreline of southern Biscayne Bay to approximate conditions in the coastal wetlands and bay that existed prior to construction of canals and water control structures. An increase in pink shrimp (*Farfantepenaeus duorarum*) density to ≥ 2 individuals m^−2^ during the wet season (i.e., August-October) along the mainland shoreline was previously proposed as an indicator of BBCW success. This study examined pre-BBCW baseline densities and compared them with the proposed target. Densities were monitored by seasonal (wet, dry) throw-trapping (1 m^2^ replicated in triplicate) at 47 sites along ~22 km of the southwestern Biscayne Bay coastline over 10 years (2007-2016). Densities varied across years and were most often higher in dry seasons. Quantile regression revealed density limitation by four habitat attributes: water temperature (°C), depth (m), salinity (ppt), and submerged aquatic vegetation (SAV: % cover). Procrustean analyses that tested for congruence between shrimp densities and habitat metrics found that water temperature, water depth, and salinity explained ~ 28%, 28%, and 22% of density variability, respectively. No significant relationship with SAV was observed. Hierarchical clustering was used to identify spatially and temporally similar groupings of pink shrimp densities by sites or season-years. Significant groupings were later investigated with respect to potentially limiting habitat attributes. Six site and four year-season clusters were identified. Although habitat attributes significantly differed among spatial clusters, within-cluster median pink shrimp densities did not correlate with within-cluster minima, maxima, medians, or standard deviations of habitat attributes. Pink shrimp densities corresponded significantly with salinity and appeared limited by it. Salinity is an environmental attribute that will be directly influenced by CERP implementation.

## Introduction

Biscayne Bay is a coastal lagoon located adjacent to the city of Miami, Florida, USA. Its watershed was heavily modified during the 20^th^ century and is currently highly managed to prevent urban, suburban, and agricultural flooding while also meeting agricultural, commercial, and residential freshwater demands. The Comprehensive Everglades Restoration Plan (CERP) seeks to restore the quality, quantity, timing, and distribution of freshwater deliveries to southern Florida nearshore areas [1], including Biscayne Bay. The Biscayne Bay Coastal Wetlands (BBCW) project, a CERP component, aims to restore a more natural hydrology and salinity regime along the western bay’s southwestern shoreline[2,3]. Three actions are needed to make this improvement: (1) increasing the total volume of freshwater deliveries; (2) diverting part of point-source freshwater discharge (i.e., canal discharges) to reestablish water delivery as overland sheet flow; and (3) altering the present timing of deliveries by lengthening discharges through the wet season (May-October) and into the dry season (November-April) [3,4]. The REstoration COordination and VERification (RECOVER) team established Interim Goals (IGs) to link ecological indicator metrics to CERP activities and thus evaluate restoration performance and realization of post-implementation ecological benefits at 5-yr intervals [1].

The pink shrimp *Farfantepenaeus duorarum* is one of many ecological indicators selected to assess ecological impacts of CERP implementation [1,5]. Pink shrimp was selected to assess estuarine ecosystems due to previously suggested abundance linkages to salinity condition [1,5]. As reviewed by Zink et al. [6], salinities within polyhaline (18 ‑ 30 ppt: [7]) and euhaline (30 – 40 ppt) ranges would directly improve pink shrimp productivity. Expansion of southwestern Biscayne Bay estuarine habitat was anticipated to benefit pink shrimp residing there [8]. Indirectly, reduced stress on seagrass communities from extreme salinity fluctuations would yield increased pink shrimp abundance due to increased seagrass cover and spatial extent [5,8]. Higher abundance of pink shrimp has been reported in areas exhibiting higher and more stable salinities [8–10]; these same coastline stretches exhibit more continuous seagrass cover [11]. Areal expansion of shoal grass (*Halodule wrightii*) cover could further amplify pink shrimp abundance due to an apparent affinity for this seagrass species [12]. The stated pink shrimp IG for southwestern Biscayne Bay is “2 shrimp m^−2^ in nearshore optimal habitat (i.e., seagrasses)” during August-October peak abundance periods [1]. This IG was based upon a peak density of ~1.8 shrimp m^−2^ observed in September during a 2 yr pilot study [8].

Historically, most freshwater delivery to Biscayne Bay was through transverse glades, broad natural channels through the Miami Coastal Ridge that allowed Everglades Basin surface water drainage [13,14] and groundwater seepage [15–20]. These natural drainage features fed fresh water from the Everglades through the coastal ridge via transverse glades into creek networks that spread surface water flows along the bay’s shoreline. Canalization converted the freshwater delivery system to one dominated by pulsed point-source (i.e., canal mouth) discharges that altered benthic submerged aquatic vegetation (SAV), infaunal, epifaunal, and nekton communities [11,21–26] and lowered the water table, reducing groundwater seepage [19,20].

Post-BBCW salinity goals for southwestern Biscayne Bay (Shoal Point to Turkey Point: Fig. 1) provide oligohaline (0.5-5 ppt) and mesohaline (5-18 ppt) regimes at the shoreline trending towards 20 ppt (polyhaline, 18-30 ppt) 500 m from the coast [4]. These conditions are anticipated to enrich estuarine faunal assemblages as well as increase estuarine species distributions and abundances [4,27,28]. Expansion of continuous submerged aquatic vegetation (SAV) habitats dominated by *Halodule wrightii*, a species commonly associated with low and variable salinity, is foreseen [11,24–26,29,30]. BBCW implementation goals for benthic habitat include increased spatial extent of nearshore seagrass beds, especially seaward expansion of *H. wrightii* [30]. Increased overlap of optimal salinity conditions with preferred benthic SAV habitats would yield indirect, synergistic benefits to estuarine fauna such as pink shrimp [4,5,27,31].

**Fig. 1.**
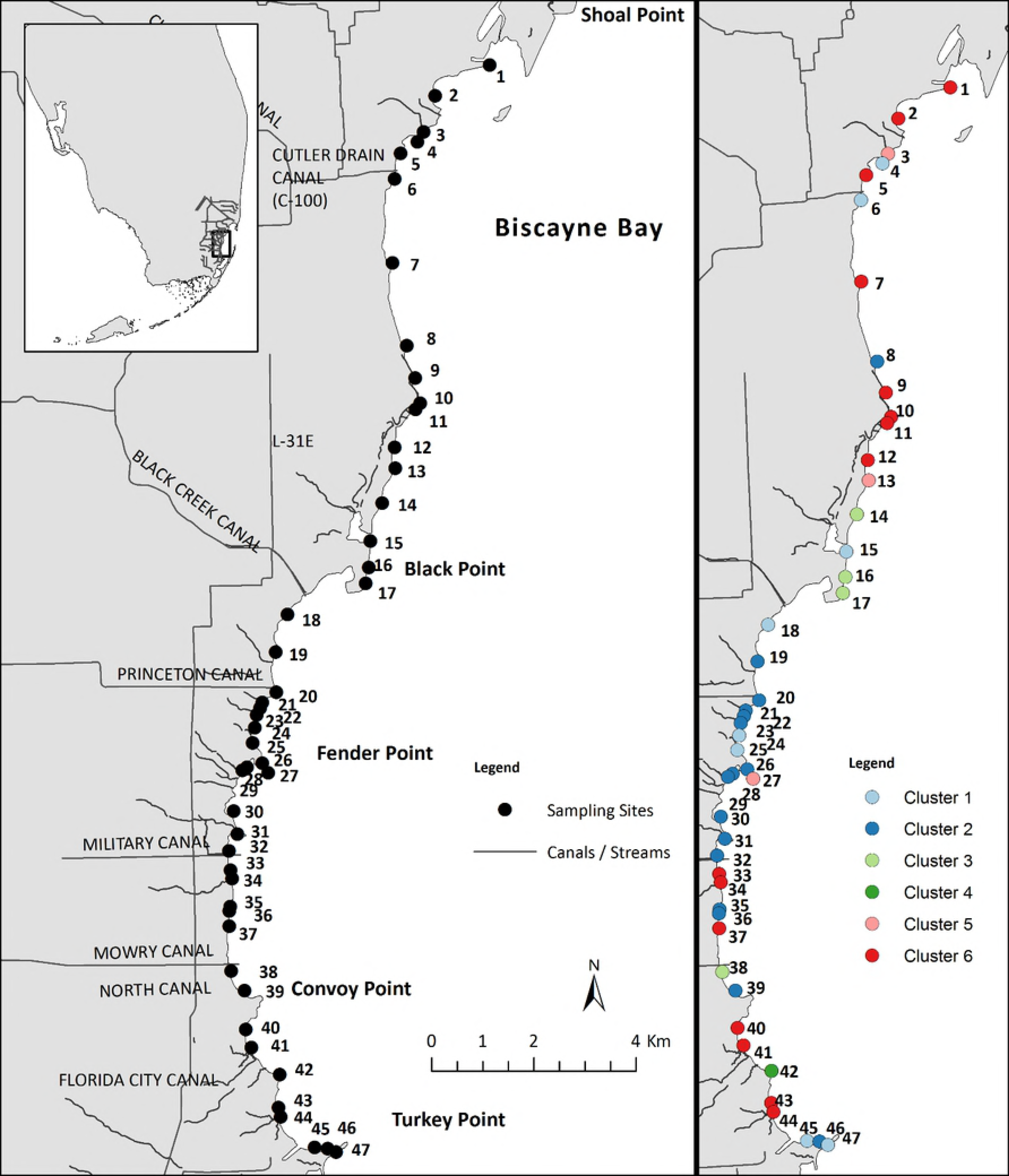
Map of study area, including referenced geographical features, and location of survey sites. The second panel depicts the same sites color-coded to match significant site clusters (Fig. 3.3B and 3.4).

The purpose of this study was to investigate spatiotemporal trends in pink shrimp density along the southwestern Biscayne Bay shoreline. We investigated the plausibility of the post-CERP establishment of ≥2 shrimp m^−2^ IG. Further, we address presumptions that (1) pink shrimp peak abundance occurs during the wet season; and (2) nearshore mesohaline salinity goals would yield increased pink shrimp abundance in the nearshore zone. Pink shrimp density relationships to species-specific and total benthic SAV % cover, as well as SAV canopy height, were also investigated. Our focus was on evaluating temporal (i.e., seasonal and inter-annual) and spatial pink shrimp density trends relative to habitat attributes. This was achieved by (1) using quantile regression to identify habitat attributes that potentially limit pink shrimp density, (2) organizing pink shrimp density and habitat observations via heatmaps to visually assess spatiotemporal variability and trends, (3) using Procrustean analysis to measure concordance between density and habitat attribute matrices, (4) employing hierarchical clustering analysis to identify spatiotemporal density clusters, and (5) investigating distributional aspects (median, minimum, maximum, and standard deviation) of habitat attribute values (temperature, salinity, water depth, and SAV % cover) within density clusters to link density patterns to the environment. These analyses employ data from wet and dry seasons of 10 years of epifaunal community monitoring data from 47 sites within 50 m of shore spanning ~22 km of shoreline.

## Materials and methods

### Study area

Biscayne Bay is a large (1,110 km^2^), shallow (depths generally < 3 m), subtropical lagoon system extending approximately 56 km north to south along the southeast coast of Florida, USA (Fig 1). Where coastal urban development is low, the bay’s mainland and shoreline consists of mangrove-seagrass ecotone punctuated by natural tidal creeks, constructed channels, and freshwater canals [32]. Overland freshwater discharges and groundwater seepage, create a salinity gradient perpendicular to the shoreline with three salinity zones: (1) western nearshore areas usually affording the lowest salinities; (2) the bay central axis marked by near oceanic salinities; and (3) oceanic salinities near the eastern passes through barrier islands to the open ocean [24,29,33]. Tidal ranges are generally 0.5 to 1 m [34,35].

### Field surveys

Epifaunal communities and SAV habitats were surveyed seasonally at fixed sampling sites (n = 47) along the southwestern Biscayne Bay nearshore zone (0-50 m) from Shoal Point to Turkey Point (Fig. 1). Surveys were conducted in public waters under authority of Biscayne National Park (Study #: BISC 06016, Permit #: BISC-2017-SCI0022). Surveys were conducted in dry (January-March sampling) and wet (July-September sampling) seasons that characterize south Florida’s climate. The primary sampling unit was the 20 m buffer around GPS coordinates that identified permanent sampling sites for continuous (15-min) recording of salinity, conductivity, and temperature. Sites were located in the shallow, open water along the western shoreline mangrove-seagrass ecotone, an area likely to be directly affected by CERP implementation. During each survey, the 47 fixed sampling sites were visited within 3 hr of high tide over 5 to 7 days within a couple of weeks’ time or less. Water quality and habitat parameters, including water temperature (°C), salinity (ppt), pH, dissolved oxygen saturation (%), dissolved oxygen concentration (mg L^−1^), water depth (m), and sediment depth (m), were recorded at each site. Benthic habitats were assessed for species-specific SAV % cover by visual assessment of 10 replicate 0.5 m^2^ quadrats per site [24,25]. In addition, canopy height (maximum seagrass blade length) was measured to provide a topography metric. Species-specific and total SAV % cover data following the methods of Lirman et al. [25] were obtained for the period 2008 to 2016.

Epifaunal communities were sub-sampled at each site (n = 3) using an open-ended, rigid-sided aluminum box (i.e., throw trap) measuring 45 cm by 1 m^2^ [36,37]. Two 3-mm stretch-mesh cover nets affixed to opposite sides of the throw-trap upper surface prevented epifauna escape during deployment. Once deployed, the throw-trap was cleared of trapped epifauna by sweeping (n = 4) its interior from alternating directions with a metal-framed seine fitted with 3 mm stretch-mesh, while gently tapping the substrate with the seine frame. Organisms collected from each sub-sample throw-trap deployment were bagged and numbered separately for storing and processing. Samples were frozen during storage until processing. No protected species were sampled.

### Epifauna identification and measurement

Taxonomic identifications and size measurements were conducted in the laboratory. Organisms collected from each replicate throw-trap deployment at a site were maintained and processed independently of each other. Where possible, carapace length (CL, mm) and total length (TL, mm) were recorded for each farfantepenaeid shrimp. Shrimps >8.0 mm CL were identified to species primarily using petasma and thelycum (i.e., sexual) morphology, although other characteristics were also used [38–41]. Shrimps <8.0 mm CL were identified to genus due to low degree of sexual morphological development [39].

### Statistical analysis

All statistical analyses were performed using the R statistical package (The R Foundation, https://www.r-project.org/). Statistical analyses were performed with a Type 1 error criterion of α = 0.10 to reduce potential Type 2 errors. Combining the data for all three trap samples for each site, density was calculated as the sum of observed shrimps per 3 m^2^ per site. Density data were natural logarithm (x + 1) transformed before analysis to reduce influence of outlying observations.

### Potential habitat limitations on pink shrimp density

As a statistical interpretation of the ecological concept of Leibig’s Law of the Minimum [42,43], quantile regression (QR) has been presented as a method to identify species distribution or abundance limitation by specific habitat attributes by focusing specifically on the upper bound of the abundance vs. habitat attribute relationship [44–47]. Pink shrimp density was first plotted against individual habitat factors to graphically assess potential limiting factors. QRs (function ‘rq’ of package ‘quantreg’) fit to the 0.5 and 0.9 density percentiles were used to statistically identify a subset of habitat attributes that suggested limitation at the median and upper edge of the density distribution. Analyses considered water temperature (°C), salinity (ppt), pH, dissolved oxygen saturation (%), dissolved oxygen concentration (mg/L), water depth (m), sediment depth (m), and the following SAV metrics: *Thalassia testudinum* % cover, *H. wrightii* % cover, total seagrass % cover, total SAV % cover, and total SAV canopy height. As in previous studies in the same region [24–26], S*yringodium filiforme* was rarely encountered (n = 10, 1.6% of total samples) and thus was not further considered.

Multiple QR functional response shapes were investigated including linear, quadratic, cubic, log-linear, natural cubic splines (function ‘ns’ of package ‘splines’) [48], and additive quantile smoothing spline (AQSS) response curves (functions ‘rqss’ and ‘qss’ of package ‘quantreg’ [49,50]). Natural cubic splines were constructed with 3 (0.25, 0.50, and 0.75 quantiles of the predictor), 2 (0.33 and 0.66 quantiles of the predictor) and 1 (0.5 quantile of the predictor) internal knots [48]. QR coefficient confidence intervals were constructed and tested for significance by *xy*-pair bootstrapping (function ‘summary.rq’ of package ‘quantreg’). Natural cubic spline QRs were modeled without intercepts; these QRs were considered significant if each individual spline describing sub-ranges of the data was significant.

### Spatiotemporal relationships

Heatmaps were generated to visualize spatiotemporal trends in pink shrimp density and the habitat attributes found by QR to potentially limit pink shrimp density. Observation data were converted to 47 row by 20 column matrices to display their spatial (47 sampling sites) and temporal (10 yr by 2 seasons) patterns, and color gradients were used to represent the magnitude of density.

Procrustean analyses allowed direct testing of statistical concordance between matrices of shrimp density and habitat attributes [51–53]. Procrustean analysis minimizes the residual sum of squares between a target matrix (**X**: here, shrimp density) and a second, fitted matrix (**Y**: here, habitat attributes), superimposed on it by scaling, rotating, and dilating [51,52]. The Procrustean Sum of Squares (PSS, also known as Gower’s Statistic: m^2^**X**,**Y**) represents the minimized residual sum of squares from the fitting procedure and is used to assess Procrustean fit ranging from 0 to 1, with higher values presenting poorer fit [51–53]. The PSS metric is equivalent to 1 −*r*^2^, where *r* is a Pearson correlation coefficient [52]. Because the method hinges on one-to-one relationships between the matrices being compared, Procrustean analysis [54,55] cannot handle missing values. Following Adams et al. [54] and Arbour and Brown [55], missing habitat attribute values were imputed with linear regressions that included site, season, and year as potential factors. PROTEST (function ‘protest,’ package ‘vegan’, permutation n = 9999) provided statistical significance of Procrustean fits between density and habitat attribute matrices [51].

Hierarchical clustering procedures were used to identify groups with similar density among sites or year-seasons [56,57]. Bray-Curtis dissimilarity matrices were constructed (‘vegdist’ function, ‘vegan’ package) with respect to site (i.e., spatial) and year-season (i.e., temporal) density observations. Hierarchical agglomerative clustering (function ‘hclust’) using the “Ward.D2” agglomeration method identified spatially and temporally similar density groupings. *A prioiri* statistical significance of clusters was tested via similarity profiling (function ‘simprof’ of package ‘clustsig’) (permutations = 999, number of expected groups = 1000) of identified density cluster memberships [57]. Permutational multivariate ANOVA (PERMANOVA: function ‘adonis2’ of package ‘vegan’) testing provided *a posteriori* cluster significance [58,59]. PERMANOVA was also used to investigate inter-annual and seasonal density differences using year-season cluster membership as a categorical nesting factor. To investigate potential dispersion influences on PERMANOVA significance, multivariate homogeneity of dispersions analysis (function ‘betadisper’ of package ‘vegan’) was used to test for inter-cluster differences in dispersion (i.e., distance to centroid) [60]. The density heat map was rearranged to visualize site and year-season cluster memberships.

### Pink shrimp density and habitat attributes among density clusters

Pink shrimp density and habitat attributes previously detected as potentially limiting to pink shrimp density via QR were investigated among site and year-season clusters. First, medians (± CI) of density and habitat attributes were computed for each site and year-season cluster. Confidence intervals (CIs) about median values were computed as:

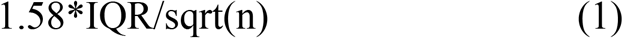

where IQR = interquantile ranges and n = sample size, as described in McGill et al. [61] and Chambers et al. [62]. Plots of density and habitat attributes’ median, CIs, minimum, and maximum values were used to visualize their distributions within site and year-season clusters. Density and habitat attributes were analyzed with respect to site or year-season clusters. Nonparametric tests were used because parametric normality and equality of variance assumptions were usually violated. Kruskal-Wallis tests were used to investigate differences in distribution shape and range (i.e., location: [63]) of density and habitat attributes among site or year-season clusters. Post-hoc Tukey-type nonparametric Conover multiple comparison tests (function ‘posthoc.tukey.conover.test’ of package ‘PMCMR’) were used to test for significant pairwise differences. These tests were implemented as χ^2^ distributions to correct for data ties, and p-values were Bonferroni-corrected [63,64]. A series of correlation analyses was used to identify habitat attribute distribution characteristics that associated with site or year-season cluster median densities. Pearson correlation analyses were applied to median, minimum, maximum, and standard deviation of habitat attributes within site or year-season clusters.

## Results

A total of 3,179 penaeid shrimp specimens were collected. The distribution of shrimp sizes suggested a gear capture inefficiency for individuals <5mm CL; therefore, data only for shrimps ≥5 mm CL (2,417 shrimps) were retained for further analysis (Fig. S1). Of the retained shrimp, 1,573 (65.1%) were identified as *F. duorarum* and the remaining 844 (34.9%) were identified as farfantepenaeids due to difficulties with species identification of individuals <8 mm CL. Of the 1,937 individuals with measured CL, 1,931 individuals (79.9%) were considered juveniles (≤17.5 mm CL) and the remaining 36 individuals were subadults.

Pink shrimp density observations ranged from 0 to 13.0 shrimp m^−2^; 105 instances (11.2%, N = 940 samples) of densities ≥2 shrimp m^−2^ were observed, while no penaeid shrimps were observed in 377 samples (40.1%). Overall, shrimp density averaged 0.86 (SD = 1.32) shrimp m^−2^, and was significantly lower (t_(α=0.10,2),939_ = −26.53, P <0.0001) than the 2 shrimp m^−2^ CERP Interim Goal threshold. Average density in any year-season was always < 2.0 shrimp m^−2^ (Table 1), although the highest year-season density (2014 Dry: 1.62 ± 2.02 shrimp m^−2^; Table 1) was the only case that did not significantly differ from 2 shrimp m^−2^ (t_(α=0.10,2),46_ = −1.30, p > 0.10). Averaged over all sites, mean dry season shrimp densities were higher than those of the subsequent wet season 50% of the time. Averaged over year-seasons, the highest mean site density was 2.15 (±1.95) shrimp m^−2^ at site 33 (Table 2). Seven sites (7, 10, 12, 33, 34, 43, and 44: 14.9%) exhibited temporally averaged densities that did not significantly differ from 2.0 shrimp m^−2^ (t_(α=0.10,2),19_ = −0.080, −1.69, −0.30, 0.34, −1.63, −1.57, −0.32, respectively; p > 0.10). Most sites (n =33, 70.2%) exhibited average densities below 1 shrimp m^−2^ (Table 2).

**Table 1:** Number of pink shrimp collected, average pink shrimp density (± SD), and average (± SD) of water quality and habitat attributes for survey year-seasons.

| Year | Season | # Shrimp | Density (# $m^{-2}$) | Temp ($^{\circ}C$) | Sal (ppt) | Depth (m) | SAV (% Cover) |
| --- | --- | --- | --- | --- | --- | --- | --- |
| 2007 | Dry | 131 | $0.93 \pm 1.11$ | $23.90 \pm 1.99$ | $26.11 \pm 5.62$ | $0.67 \pm 0.18$ | NA |
| | Wet | 63 | $0.45 \pm 0.9$ | $30.81 \pm 1.07$ | $20.36 \pm 5.98$ | $0.72 \pm 0.17$ | NA |
| 2008 | Dry | 165 | $1.17 \pm 1.95$ | $22.29 \pm 1.44$ | $25.28 \pm 4.12$ | $0.73 \pm 0.17$ | $52.45 \pm 26.52$ |
| | Wet | 104 | $0.74 \pm 1.09$ | $29.84 \pm 0.99$ | $23.57 \pm 5.32$ | $0.73 \pm 0.17$ | $70.38 \pm 15.97$ |
| 2009 | Dry | 198 | $1.4 \pm 1.4$ | $21.61 \pm 1.53$ | $25.95 \pm 5.61$ | $0.63 \pm 0.15$ | $78.36 \pm 14.13$ |
| | Wet | 56 | $0.4 \pm 0.58$ | $31.25 \pm 1.85$ | $22.93 \pm 6.47$ | $0.65 \pm 0.15$ | $75.25 \pm 20.76$ |
| 2010 | Dry | 98 | $0.7 \pm 0.94$ | $19.28 \pm 2.89$ | $25.50 \pm 2.90$ | $0.59 \pm 0.14$ | $62.28 \pm 23.08$ |
| | Wet | 123 | $0.87 \pm 1.34$ | $31.62 \pm 1.17$ | $24.18 \pm 6.92$ | $0.71 \pm 0.20$ | $63.46 \pm 31.63$ |
| 2011 | Dry | 98 | $0.7 \pm 0.9$ | $21.31 \pm 1.83$ | $27.09 \pm 3.02$ | $0.60 \pm 0.18$ | $72.35 \pm 19.22$ |
| | Wet | 112 | $0.79 \pm 0.99$ | $31.72 \pm 1.56$ | $31.86 \pm 3.68$ | $0.73 \pm 0.22$ | $69.96 \pm 22.34$ |
| 2012 | Dry | 182 | $1.29 \pm 1.73$ | $22.52 \pm 1.00$ | $24.47 \pm 3.45$ | $0.71 \pm 0.16$ | $83.74 \pm 12.07$ |
| | Wet | 182 | $1.29 \pm 1.65$ | $31.10 \pm 1.85$ | $15.30 \pm 5.64$ | $0.73 \pm 0.15$ | $65.44 \pm 16.61$ |
| 2013 | Dry | 82 | $0.58 \pm 0.68$ | $21.45 \pm 1.07$ | $28.85 \pm 3.41$ | $0.86 \pm 0.18$ | $57.98 \pm 19.80$ |
| | Wet | 29 | $0.21 \pm 0.43$ | $29.30 \pm 0.87$ | $15.98 \pm 7.24$ | $0.79 \pm 0.18$ | $60.07 \pm 17.74$ |
| 2014 | Dry | 228 | $1.62 \pm 2.02$ | $22.89 \pm 1.39$ | $23.33 \pm 5.71$ | $0.69 \pm 0.15$ | $61.41 \pm 20.49$ |
| | Wet | 15 | $0.11 \pm 0.22$ | $29.44 \pm 1.12$ | $29.30 \pm 4.84$ | $1.02 \pm 0.20$ | $53.20 \pm 20.96$ |
| 2015 | Dry | 92 | $0.65 \pm 0.81$ | $26.06 \pm 1.48$ | $28.85 \pm 3.42$ | $0.74 \pm 0.19$ | $68.51 \pm 19.71$ |
| | Wet | 204 | $1.45 \pm 2.25$ | $30.87 \pm 1.45$ | $23.11 \pm 6.19$ | $0.80 \pm 0.17$ | $64.23 \pm 19.75$ |
| 2016 | Dry | 119 | $0.84 \pm 1.18$ | $26.03 \pm 2.06$ | $18.60 \pm 6.08$ | $0.77 \pm 0.20$ | $60.63 \pm 21.48$ |
| | Wet | 136 | $0.96 \pm 0.93$ | $30.28 \pm 1.22$ | $12.22 \pm 2.94$ | $0.83 \pm 0.17$ | $59.73 \pm 21.19$ |
| Overall | Dry | 1393 | $0.99 \pm 1.38$ | $22.73 \pm 2.65$ | $25.40 \pm 5.27$ | $0.70 \pm 0.19$ | $66.54 \pm 22.00$ |

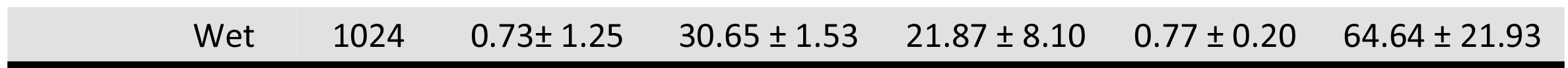

**Table 2:**
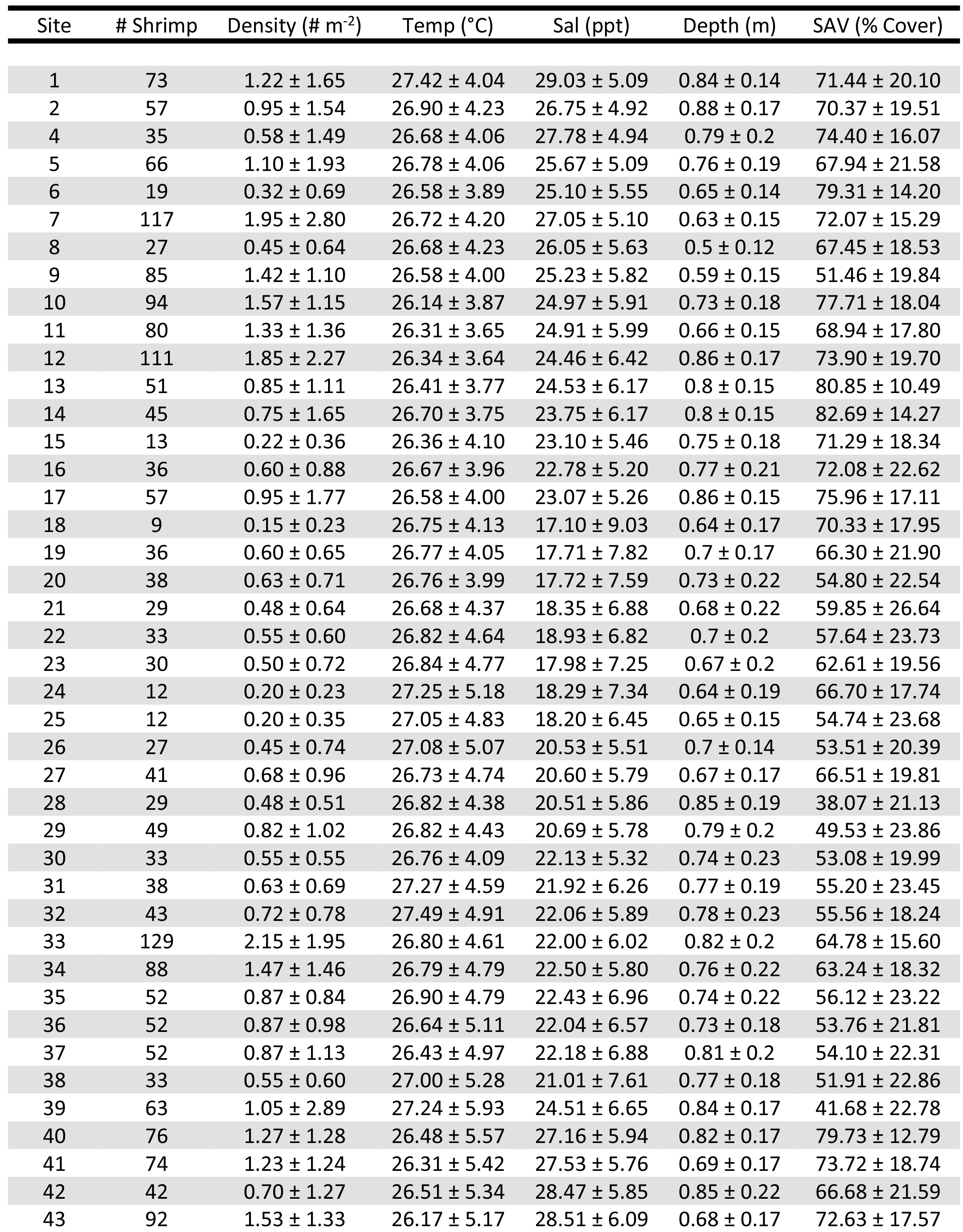
Number of pink shrimp collected, average pink shrimp density (± SD), and average (± SD) of water quality and habitat attributes for survey sites.

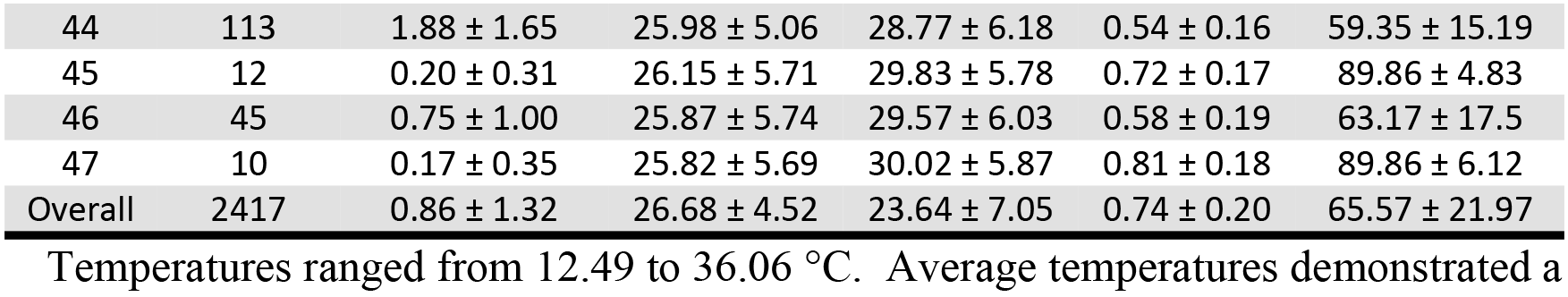

Temperatures ranged from 12.49 to 36.06 °C. Average temperatures demonstrated a clear pattern of cooler (22.73 ± 2.65 °C) and warmer (30.65 ± 1.53 °C) values for dry and wet seasons, respectively (Table 1, Fig. 3C). The dry season record was punctuated by an extreme cold front event that occurred during the 2010 field sampling. No pattern of variation in average temperatures among sites was readily discernable (Table 2, Fig. 3C). Salinities ranged from 2.48 to 39.71 ppt; overall average salinity was 23.64 (± 7.05) ppt (Table 2). Spatially averaged wet season salinities were generally lower than those of dry seasons, although 2011, 2014, and 2015 wet seasons were notable exceptions with higher average salinity than both the preceding and following dry seasons (Table 1). Sampling sites’ mean salinity and standard deviation of salinity were negatively correlated (Pearson r = −0.63, t = −5.49, d.f. = 45, p < 0.0001; Fig. S2). Wet 2011 and 2015 were considered ‘hypersaline’ due to duration of hypersaline (>40 ppt) conditions observed in these year-seasons [65]. Only four temporally averaged site salinities were mesohaline, most (n = 42) were polyhaline, and one was euhaline (Table 2). Water depths ranged from 0.19 to 1.5 m and averaged 0.74 (± 0.20) overall (Table 2) with no appreciable trends among year-seasons or among sites (Fig. 3E). Total SAV % cover ranged from 4.57 to 100% and averaged 66.57% (± 21.97) with no clear year-season variation patterns (Table 1, 2; Fig. 3F). A planktonic microalgal bloom event was observed in parts of the Biscayne Bay coastal area during the 2013 wet season [65,66].

### Habitat limitations on pink shrimp density

Of the multiple habitat attributes investigated, significant QR analysis results revealed that temperature (°C), salinity (ppt), water depth (m), and SAV (% cover) potentially limited pink shrimp density (Table 3, Fig 2). QR of density vs. temperature yielded a single-knot natural cubic spline relationship. This relationship was roughly dome-shaped and maximized at 26.6 °C, with tails that tapered off at higher and lower temperatures (Table 3, Fig 2A). Temperatures between 21.08 and 31.33 °C did not appear to limit pink shrimp densities to <2 shrimp m^−2^ (Fig2A). Although a series of functional shapes was considered for the QR density vs. salinity response curve, only the linear and log-linear responses were found to be both significant and ecologically plausible [67]. The log-linear response, which suggested severe density limitation below10 ppt and asymptotic at salinities above 10 ppt (Fig. 2B), seemed more plausible than the linear response. Salinities < ~18 ppt limited shrimp density to <2 shrimp m^−2^ (Fig. 2B). QR of pink shrimp density against water depth (m) yielded a 3 knot (0.25, 0.5, 0.75 quantile) splined relationship with steep increases in limitation below ~0.6 m and above ~ 1.0 m and a bimodal midsection (Fig. 2C). Apparent limitation of density to <2 shrimp m^−2^ occurred at water depths less than 0.43 m and greater than 1.05 m (Fig. 2C). Shrimp density had a logarithmic linear relationship with SAV % cover (Fig. 2D). SAV cover less than 45% limited density to <2 shrimp m^−2^ (Fig. 2D). For the four habitat attributes, significant QRs were observed at the 0.9 quantile, but not at the 0.5 quantile (Table 3).

**Table 3:**
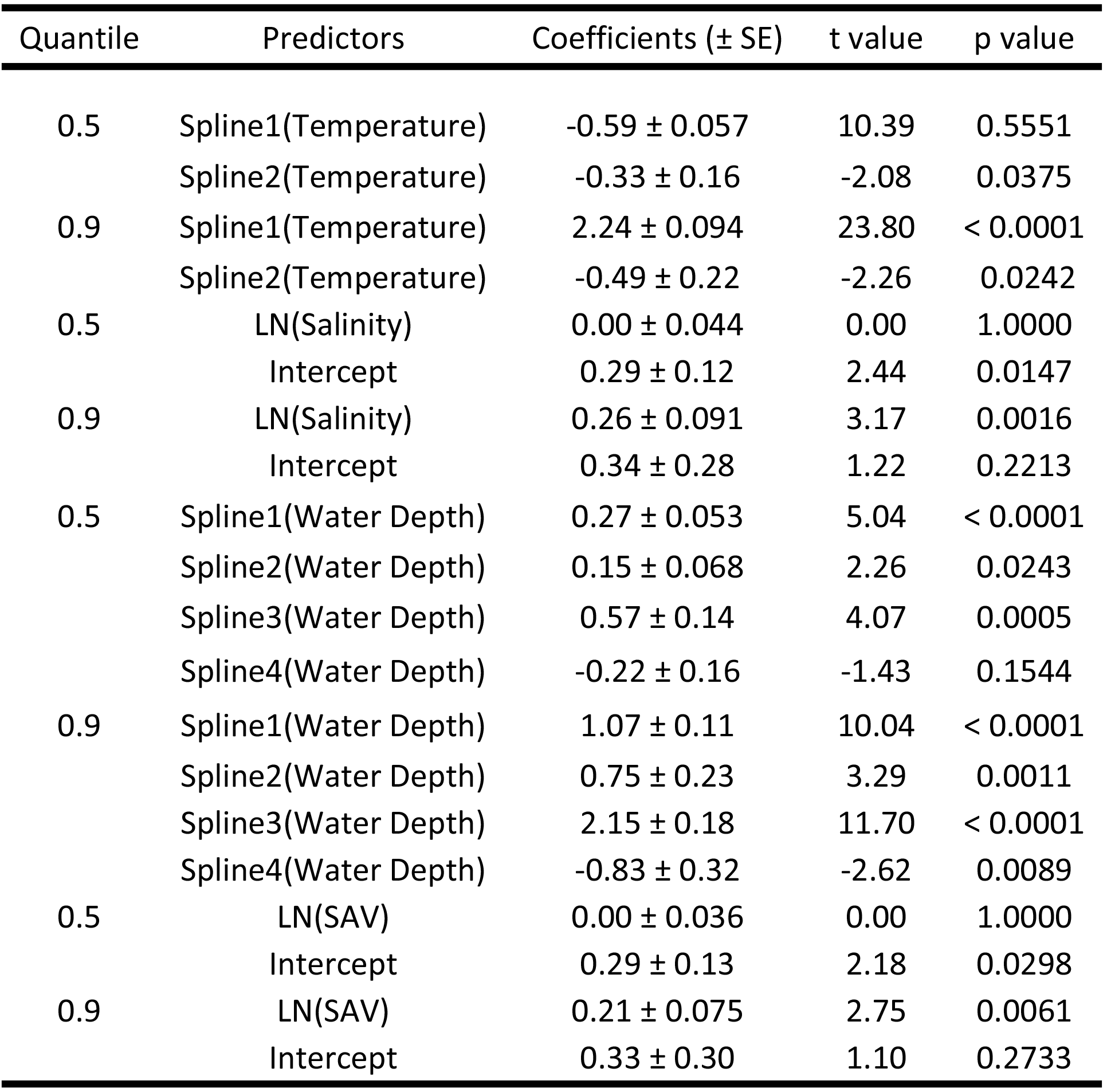
Statistical significance of 0.5 and 0.9 quantile regressions of pink shrimp density (shrimp m^−2^: LN([x+1]) against temperature (°C), salinity (ppt), water depth (m), and submerged aquatic vegetation (SAV: % cover). LN = natural logarithm.

| Quantile | Predictors | Coefficients (± SE) | t value | p value |
| --- | --- | --- | --- | --- |
| 0.5 | Spline1(Temperature) | -0.59 ± 0.057 | 10.39 | 0.5551 |
|  | Spline2(Temperature) | -0.33 ± 0.16 | -2.08 | 0.0375 |
| 0.9 | Spline1(Temperature) | 2.24 ± 0.094 | 23.80 | < 0.0001 |
|  | Spline2(Temperature) | -0.49 ± 0.22 | -2.26 | 0.0242 |
| 0.5 | LN(Salinity) | 0.00 ± 0.044 | 0.00 | 1.0000 |
|  | Intercept | 0.29 ± 0.12 | 2.44 | 0.0147 |
| 0.9 | LN(Salinity) | 0.26 ± 0.091 | 3.17 | 0.0016 |
|  | Intercept | 0.34 ± 0.28 | 1.22 | 0.2213 |
| 0.5 | Spline1(Water Depth) | 0.27 ± 0.053 | 5.04 | < 0.0001 |
|  | Spline2(Water Depth) | 0.15 ± 0.068 | 2.26 | 0.0243 |
|  | Spline3(Water Depth) | 0.57 ± 0.14 | 4.07 | 0.0005 |
|  | Spline4(Water Depth) | -0.22 ± 0.16 | -1.43 | 0.1544 |
| 0.9 | Spline1(Water Depth) | 1.07 ± 0.11 | 10.04 | < 0.0001 |
|  | Spline2(Water Depth) | 0.75 ± 0.23 | 3.29 | 0.0011 |
|  | Spline3(Water Depth) | 2.15 ± 0.18 | 11.70 | < 0.0001 |
|  | Spline4(Water Depth) | -0.83 ± 0.32 | -2.62 | 0.0089 |
| 0.5 | LN(SAV) | 0.00 ± 0.036 | 0.00 | 1.0000 |
|  | Intercept | 0.29 ± 0.13 | 2.18 | 0.0298 |
| 0.9 | LN(SAV) | 0.21 ± 0.075 | 2.75 | 0.0061 |
|  | Intercept | 0.33 ± 0.30 | 1.10 | 0.2733 |

**Fig. 2.**
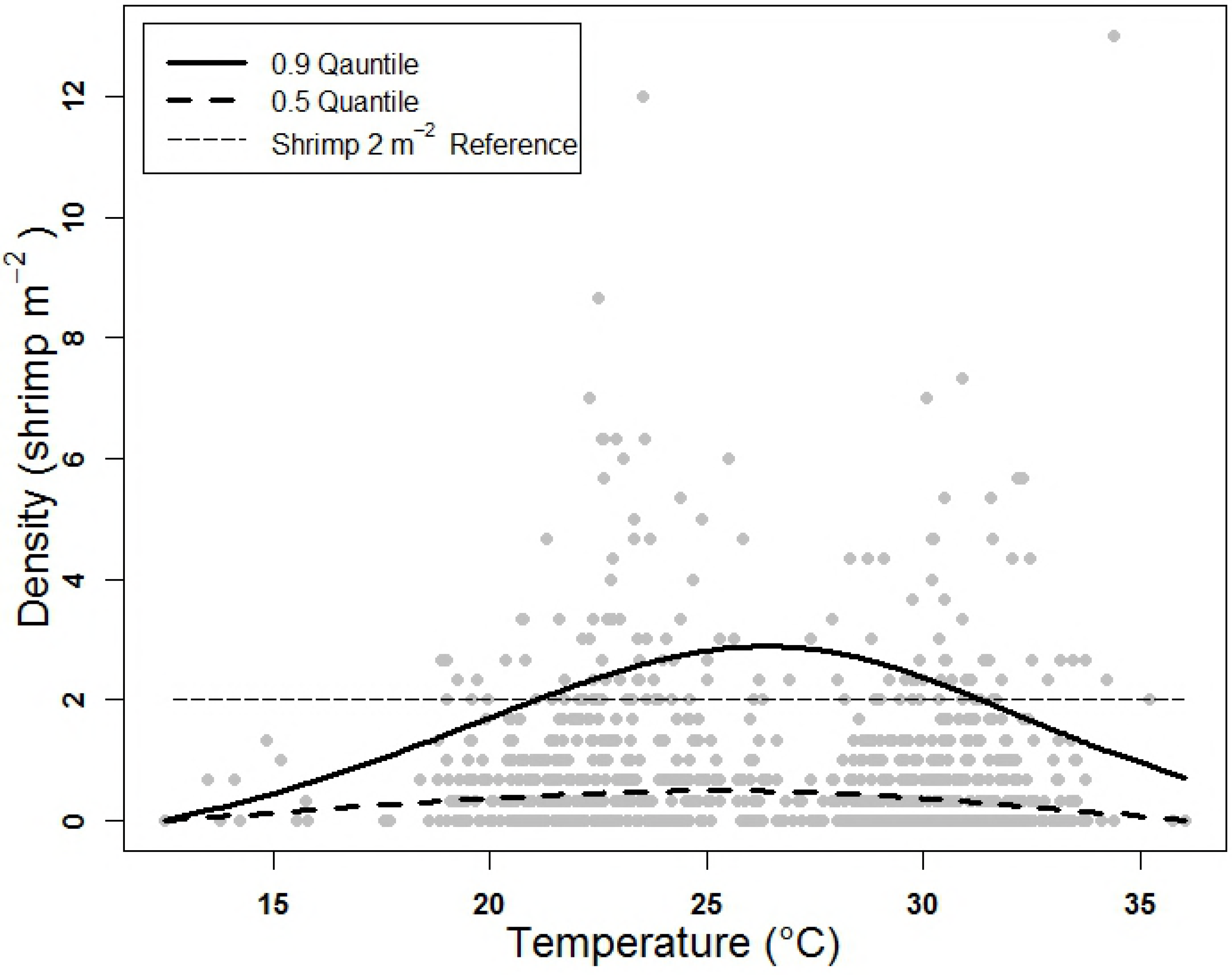
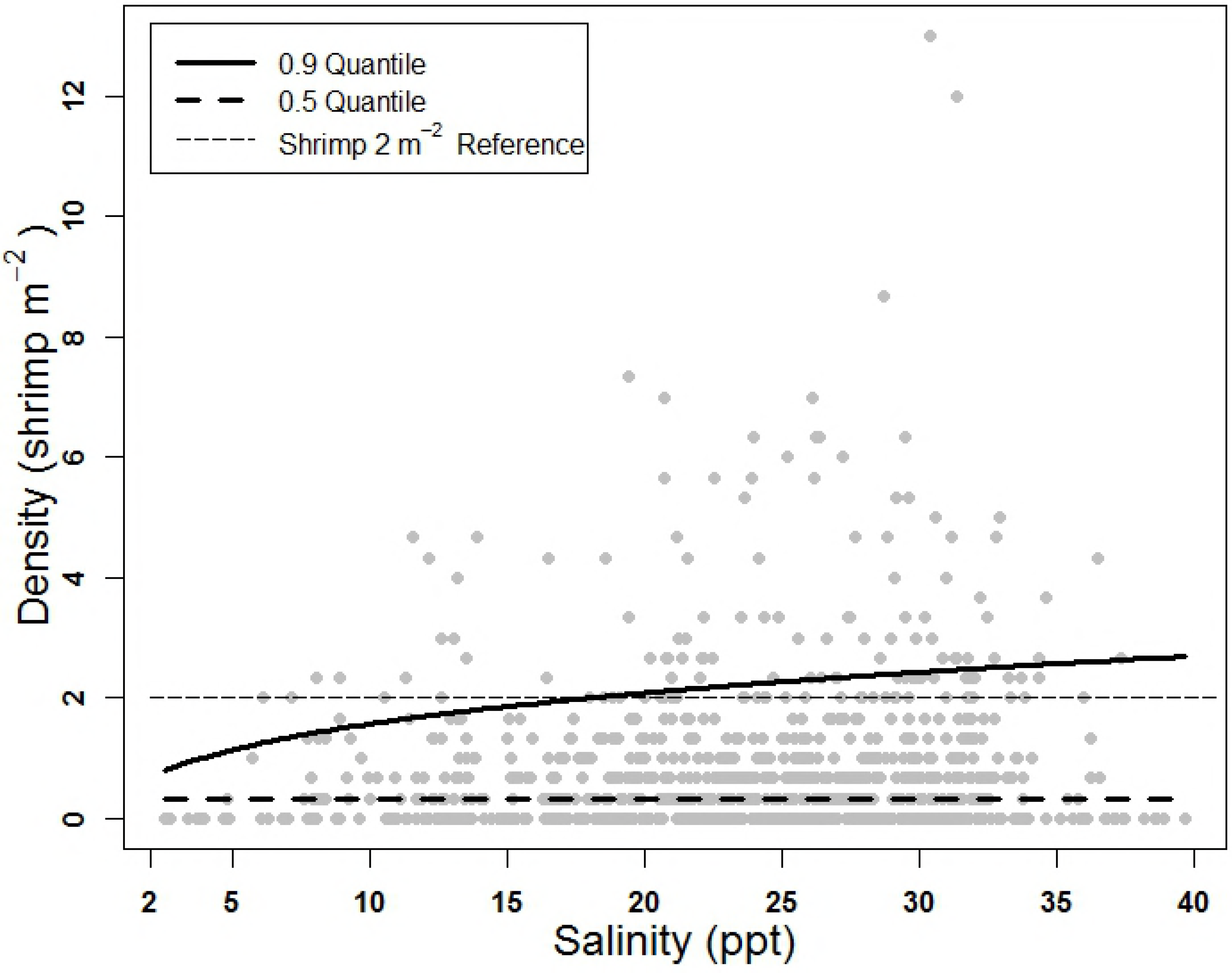
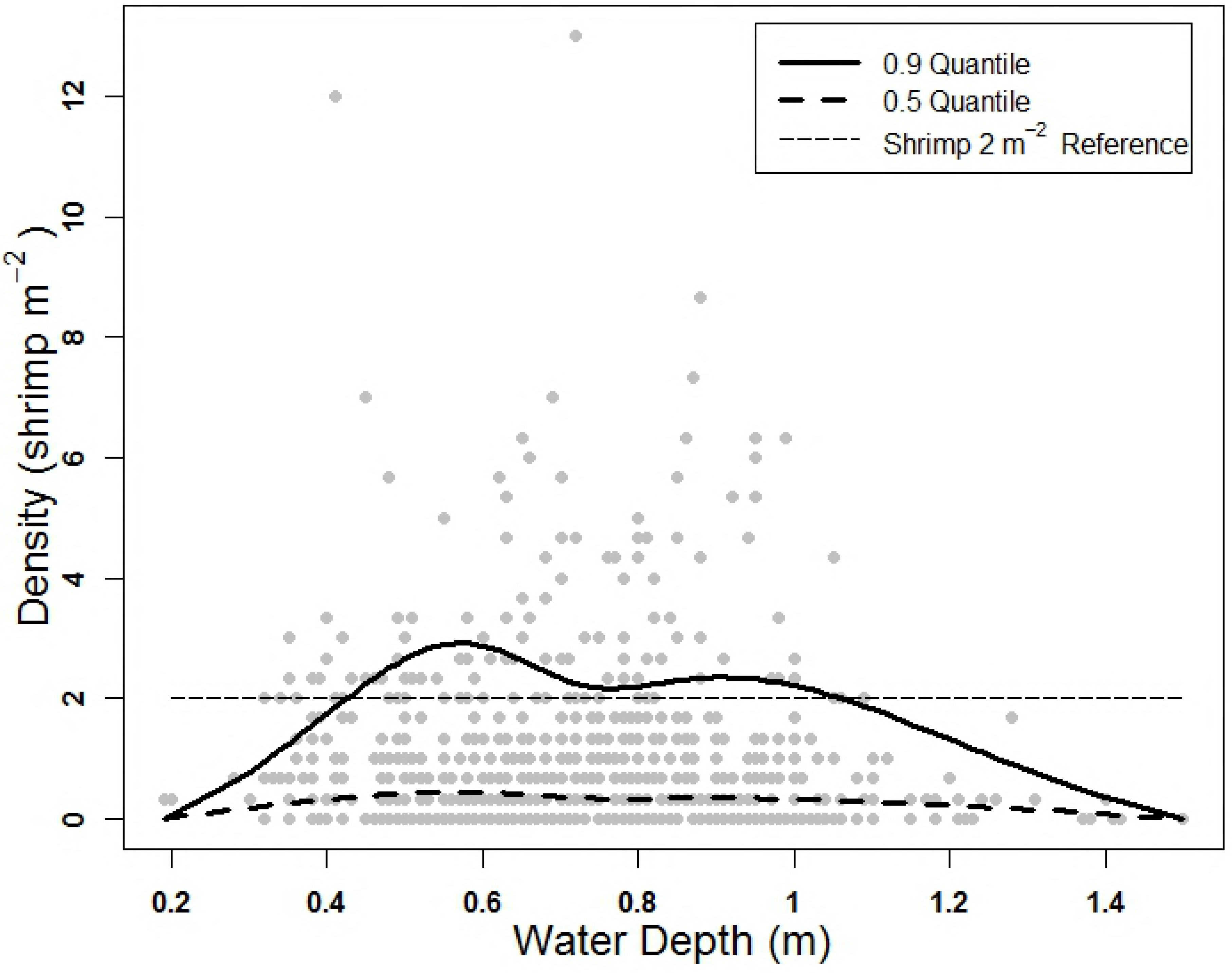
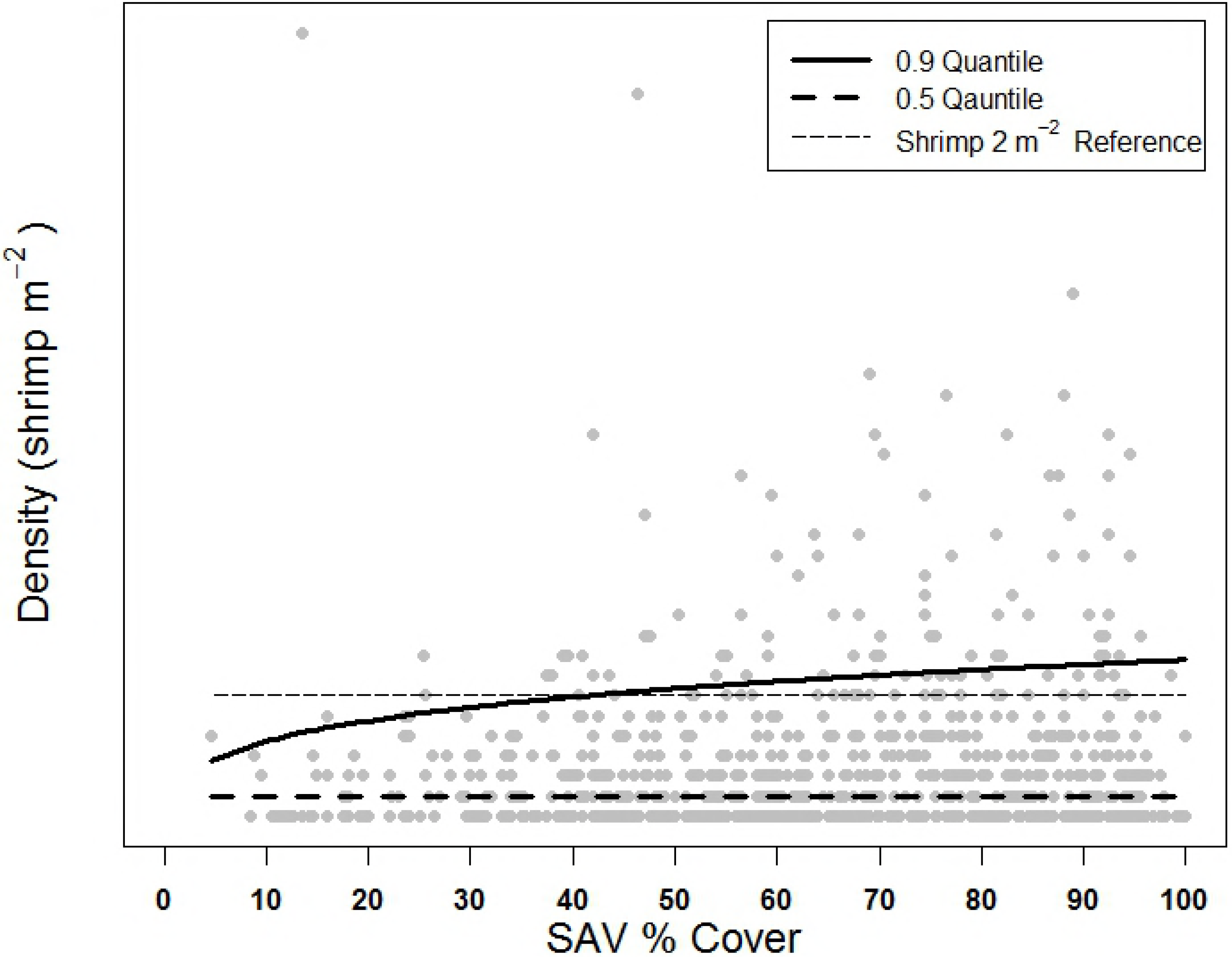
Pink shrimp density (shrimp m^−2^) and back-transformed 0.50 and 0.90 quantile regressions lines of predicted density (LN x+1) plotted against A) temperature (°C), B) salinity (ppt), C) water depth (m), and D) submerged aquatic vegetation (SAV: % cover). Predicted regression lines depict relationships reported in Table 2.

### Spatiotemporal relationships

Heatmap visualization of pink shrimp spatiotemporal density trends revealed a general absence of pink shrimp from sites 13 to 28 (approximately Black Point to Fender Point, Fig. 1) and sites 45 to 47 (near Turkey Point, Fig. 1) across all year-seasons (Fig. 3A). Within these groups of sites, only 16 (4.4%, N = 360) and 4 (6.7%, N=60) instances of pink shrimp densities >2 shrimp m^−2^ were observed respectively. Generally higher densities were observed in sites 31 through 44 and sites 1 to 12, where 42 (15%, N=280) and 40 (16.7%, N = 240) instances, respectively, of densities >2 shrimp m^−2^ were observed across all year-seasons. Densities were particularly low during 2007, 2009, 2013, and 2014 wet seasons (< 0.5 shrimp m^−2^: Table 1), when shrimp were absent from a high proportion of samples (55.3, 51.1, 70.2, and 76.6%, respectively: Fig 3A). Other year-seasons (2008 dry, 2009 dry, 2012 dry, 2012 wet, 2014 dry, and 2015 wet: Fig. 3A) exhibited high average density (> 1 shrimp m^−2^) because of a preponderance of higher density observations, which offset low and zero-catch observations from Black Point to Fender Point (Fig. 1). Average densities in these year-seasons yielded the highest year-season average densities, which were all >1 shrimp m^−2^ (Table 1).

**Fig. 3.**
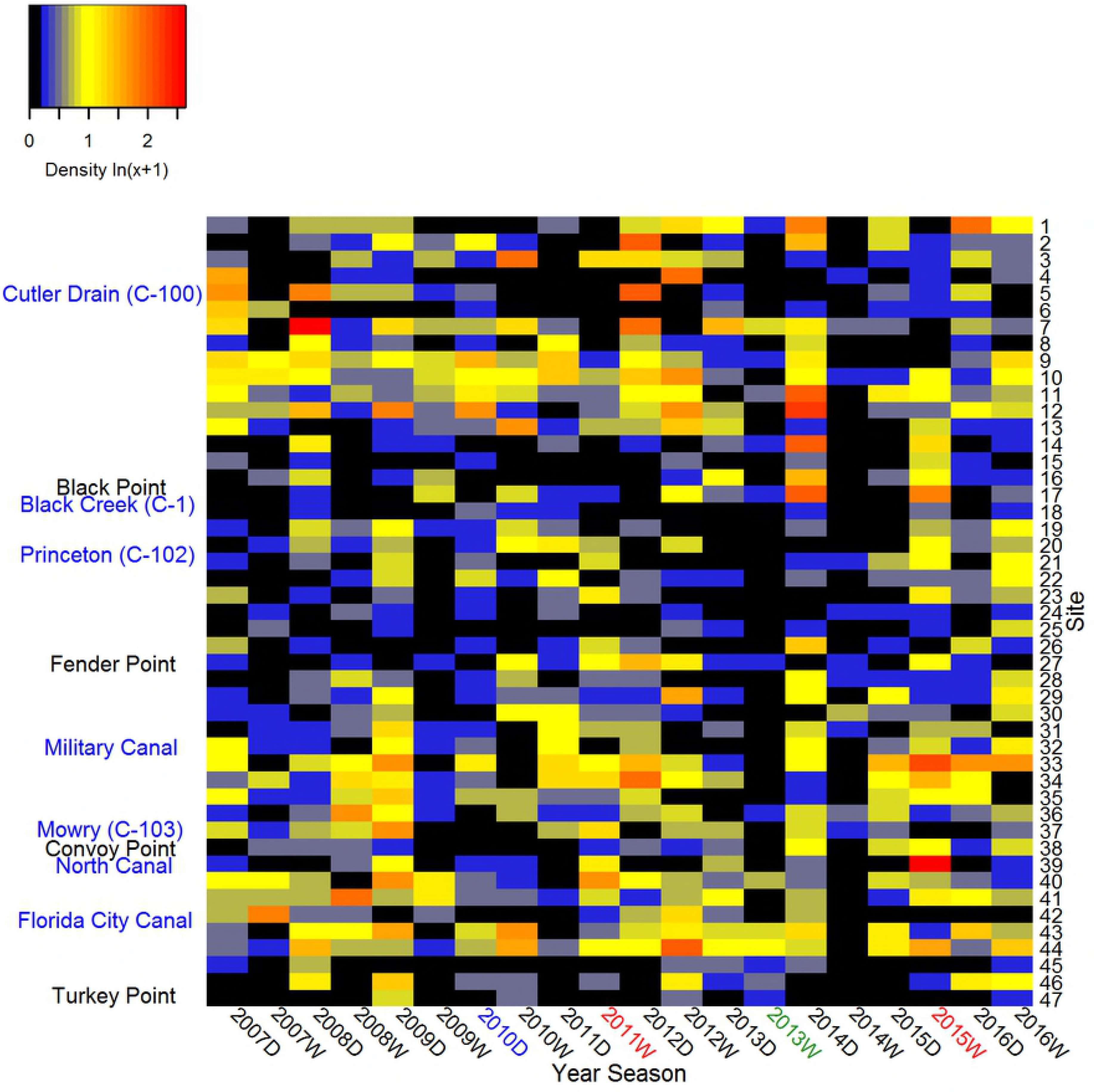
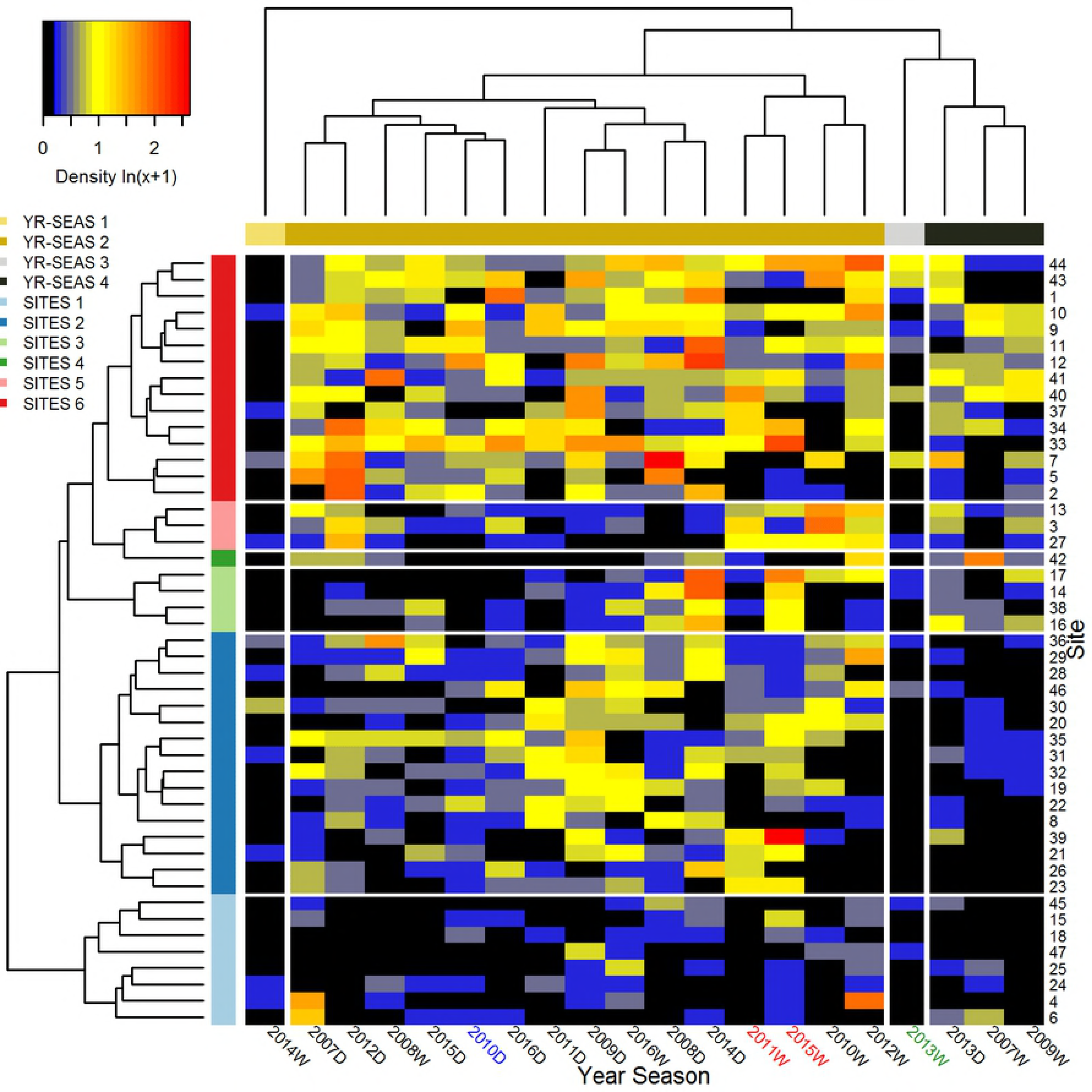
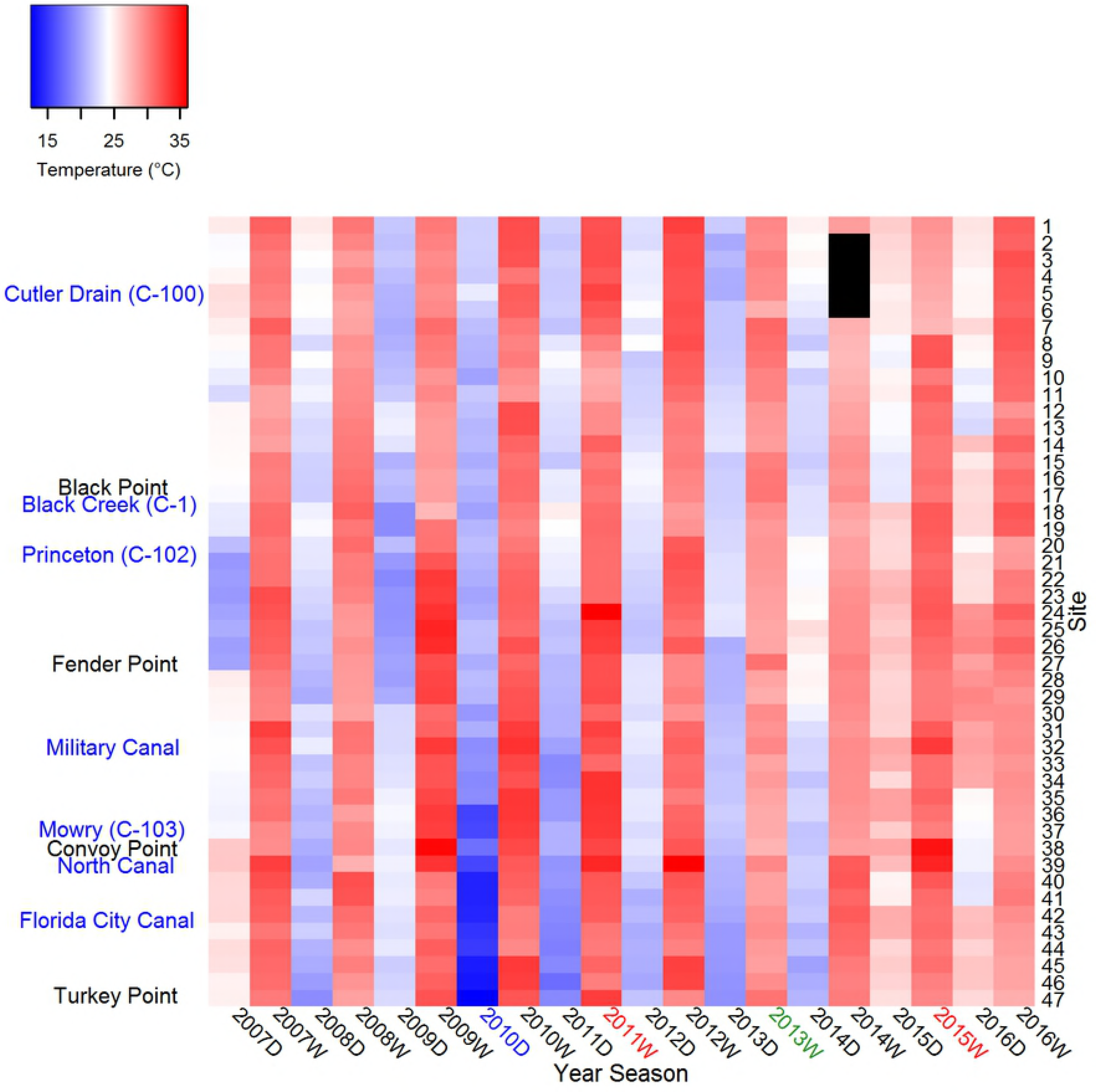
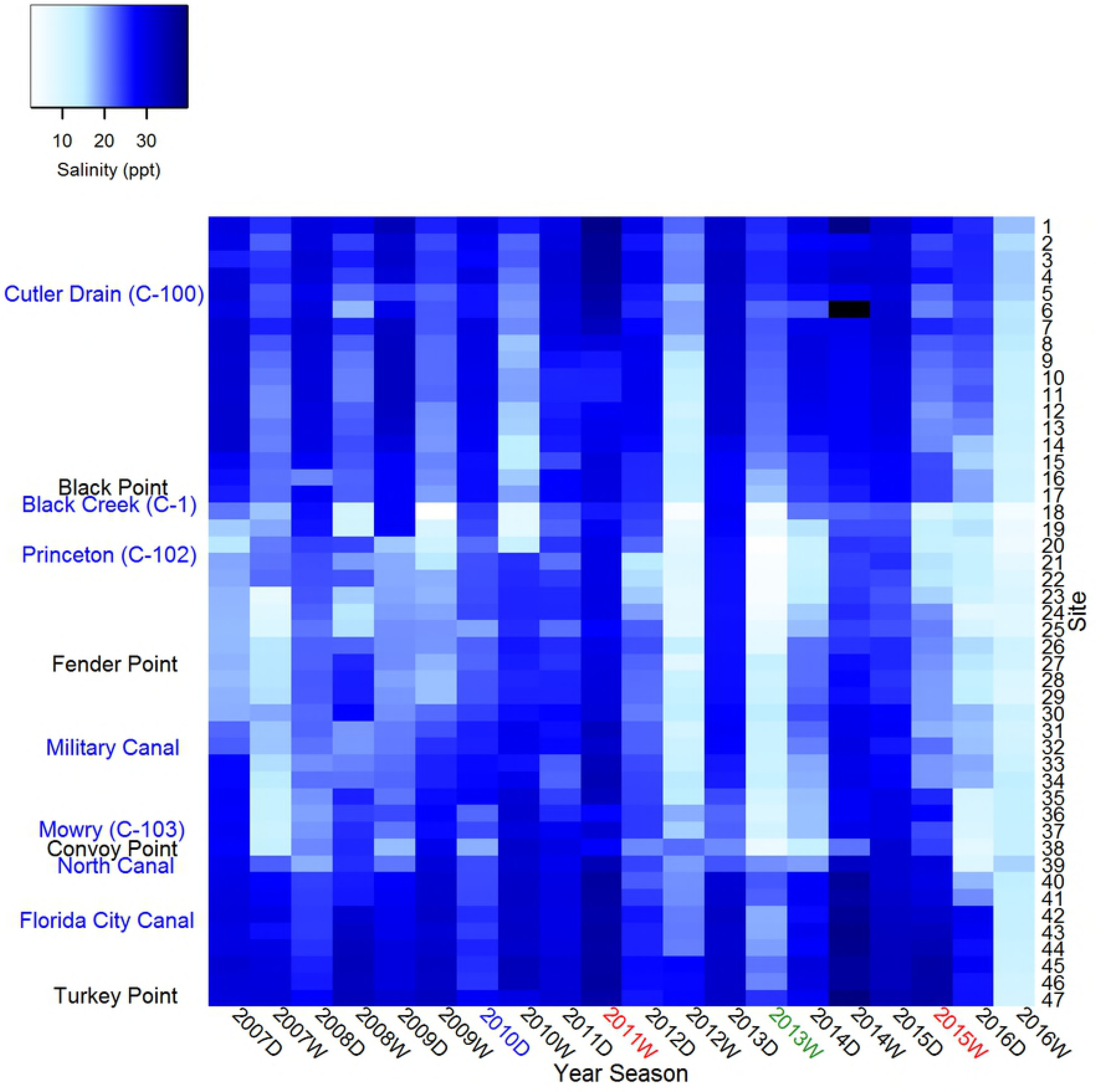
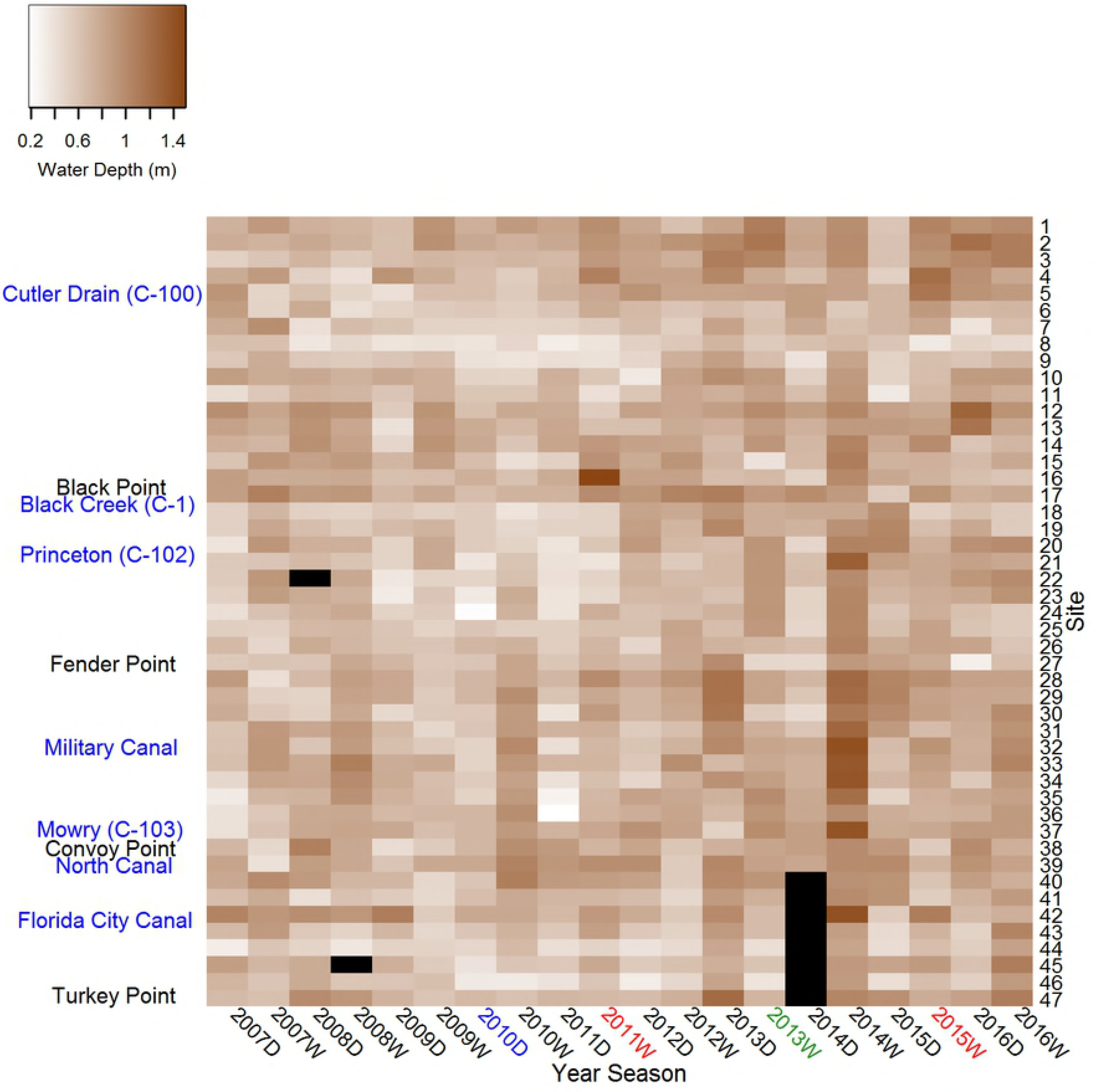
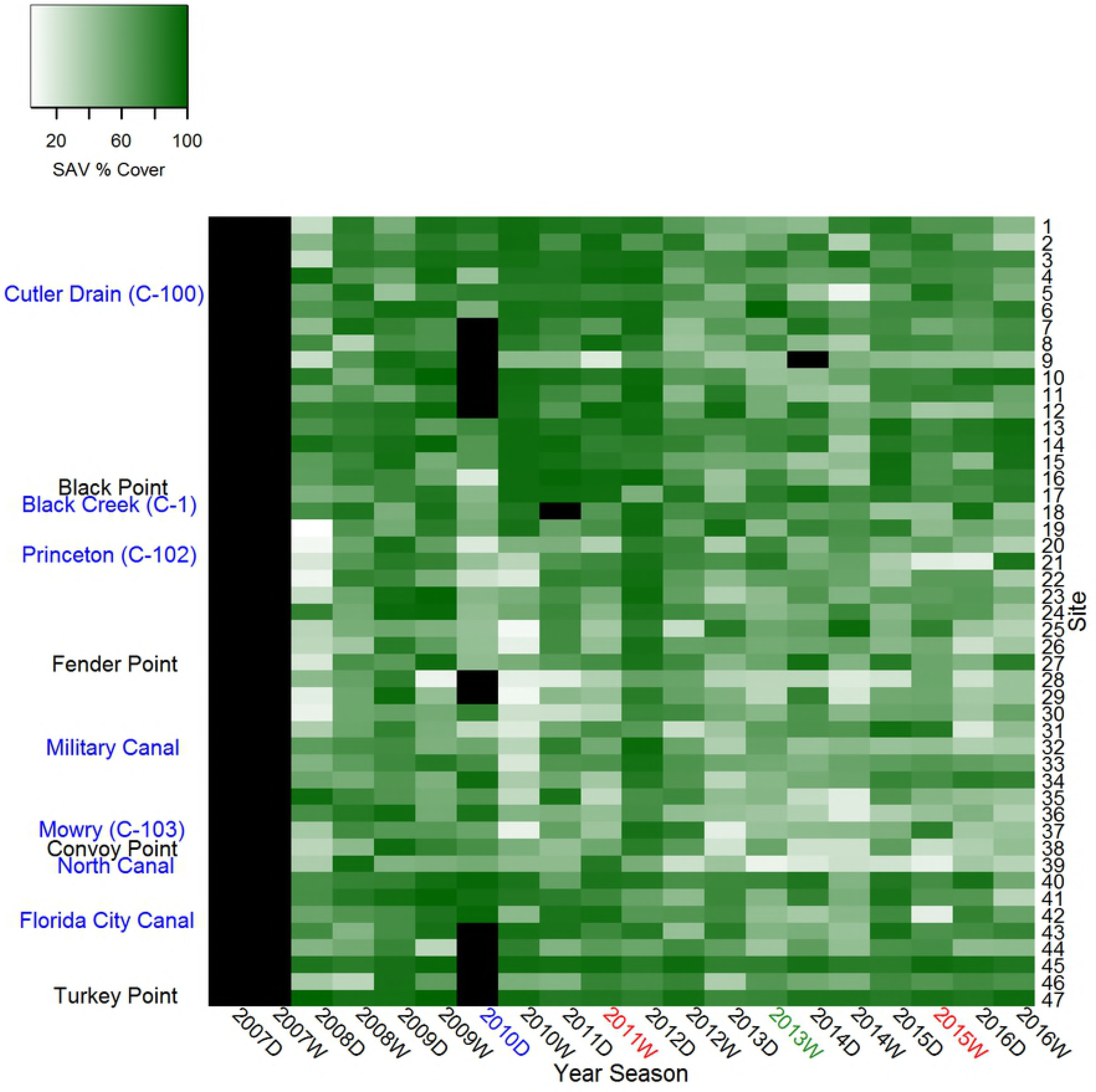
Heatmaps depicting spatial (i.e., site) and temporal (i.e., year-season) trends in A) pink shrimp density (shrimp m^−2^: LN[x+1]), C) temperature (°C), D) salinity (ppt), E) depth (m), and F) SAV (% cover). Shrimp densities are also depicted as organized (B) by site and year-season clusters. Color bars along the left and top margins of B) reflect significant sites and year-season clusters as denoted in the legend. Black cells in A) and B) highlight 0 shrimp m^−2^ observations while in C) through F) black bars represent missing values. Year-season label colors depict ecological perturbations: red = hypersalinity event, blue = cold snap, green = algal bloom. Labels on the left margin of (A) refer to canal outlets (blue) and coastline features (black) depicted in Fig. 1.

Heatmaps were also developed to visualize spatiotemporal trends in temperature, salinity, water depth, and SAV (Fig. 3.3C, D, E, F). Procrustean analyses revealed significant concordance of the shrimp density matrix (Fig. 3A) to water depth, temperature, and salinity habitat attribute matrices (Fig. 3 C, D, and E) but not to the SAV matrix (Fig. 3F, Table 4). Water depth and temperature exhibited the highest correlations, followed by salinity (Table 4). Each comparison yielded a high residual sum of squares (high m^2^**X**,**Y** values), indicating relatively weak explanatory power of individual habitat attributes (Table 4). Procrustean fitting procedures explained 28.3, 27.1, and 22.1% of the variability in density for water depth, temperature, and salinity, respectively.

**Table 4:**
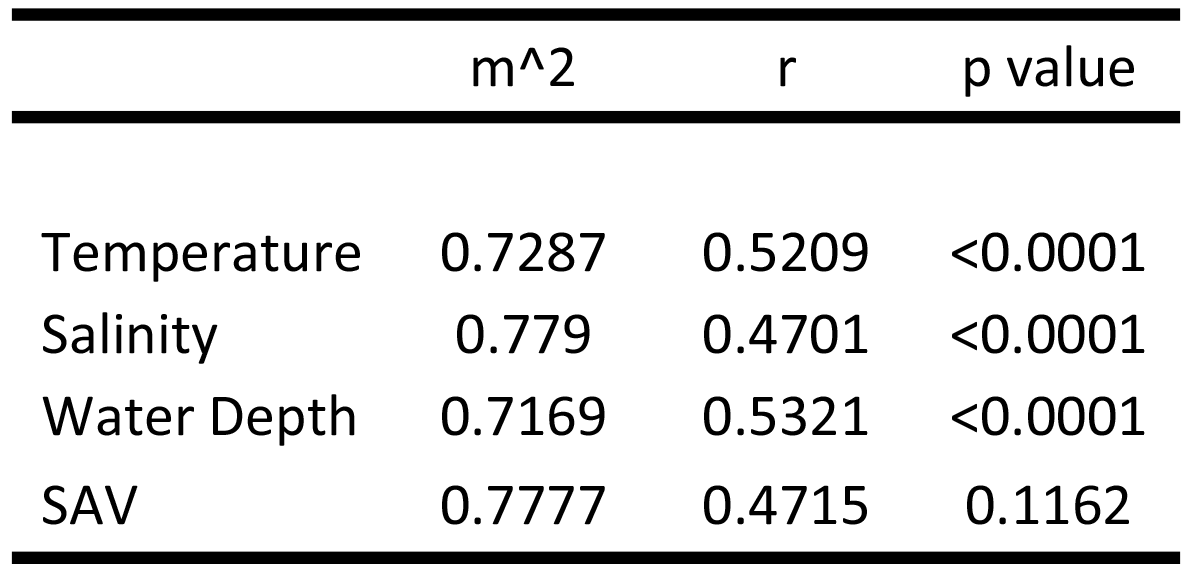
Results of Procrustean analysis of density (shrimp m^−2^: LN([x+1]) relative to temperature (°C), salinity (ppt), water depth (m), and SAV (% cover) including goodness-of-fit measure (m^2^), correlation of the Procrustean rotation (r), and p value of the fit.

SIMPROF testing identified six significant site clusters and four significant year-season clusters (Fig 3B). PERMANOVA testing of cluster membership confirmed SIMPROF site (F_5,41_ = 4.765, p = 0.001, R^2^ = 0.368) and year-season (F_7316_ = 3.727, p = 0.001, R^2^ = 0.411) clustering. Two site clusters (i.e., 2 and 6: Fig. 3B) together included most (66%, n = 31) of the sampling stations. One large year-season cluster included most year-seasons (75%, n = 15: Fig. 3B). Smaller membership year-season clusters (80%, n = 4) were mostly comprised of wet seasons representing pink shrimp densities (<0.5 shrimp m^−2^). Substantial differences in shrimp densities of members of different clusters likely drove the significantly differing multivariate dispersion among site (F_5,41_ = 14.886, p < 0.0001) and year-season (F_3,16_ = 22.987, p = <0.0001) clusters. PERMANOVA testing detected significant season (F_1,9_ = 1.912, p = 0.0063), but not year (F_9,9_ = 1.020, p = 0.4279), categorical temporal effects. Multivariate dispersions differed significantly between years (F_9,10_ = 5.56*10^29^, p < 0.0001) and seasons (F_1,18_ = 7.047, p = 0.0161), with greater observed variability in the wet season than the dry season. PERMANOVA p values were considered to be conservative because greater multivariate dispersion was observed in groups with larger sample sizes.

### Pink shrimp density and habitat attributes among density clusters

Significant differences in density distributions were detected among both site and year-season clusters (Table 5, Fig. 4A, Fig. 5A). Site clusters represented three relative median density levels: high (~0.7 shrimp m^−2^: site cluster 6); intermediate (~0.3 shrimp m^−2^: site clusters 2, 3, and 5); and low density (0.0 and ~0.14 shrimp m^−2^: site clusters 1 and 4, respectively: Table 5A, Fig. 4A). Year-season clusters also exhibited three relative density levels: high (0.51 shrimp m^−2^: year-season cluster 2), intermediate (0.29 shrimp m^−2^: year-season cluster 4) and low density (0.0 shrimp m^−2^: year-season clusters 1 and 3) (Table 5B, Fig. 5A). Significant differences were also observed for salinity, water depth, and SAV distributions among site clusters (Table 5A; Fig. 4B, C, D), while temperature distributions did not differ among site clusters. All four habitat attributes exhibited significant differences among year-season clusters (Table 5B, Fig. 5B, C, D, E). No significant correlations were found between site cluster median density and median, minimum, maximum, or standard deviation of habitat attributes within clusters. Low sample size prevented investigation of correlations of year-season clusters’ median density or maximum density with clusters’ habitat attributes’ distribution characteristics.

**Table 5:** Median and ~95% CI of density (shrimp m^−2^: LN([x+1]), temperature (°C), salinity (ppt), water depth (m), and submerged aquatic vegetation (SAV: % cover) and the χ^2^, d.f., and p values associated with Kruskal-Wallis testing of density clusters relative to A) site and B) year-season. Median CI computed as described in the text. Values in A) and B) are depicted in Fig. 4 and 5, respectively.

| Site Cluster | n | Density | Temperature | Salinity | Water Depth | SAV |
| --- | --- | --- | --- | --- | --- | --- |
| 1 | 160 | $0.00 \pm 0.036$ | $27.73 \pm 1.04$ | $21.15 \pm 1.29$ | $0.68 \pm 0.032$ | $79.00 \pm 4.00$ |
| 2 | 320 | $0.29 \pm 0.061$ | $28.50 \pm 0.71$ | $22.38 \pm 0.87$ | $0.72 \pm 0.024$ | $56.50 \pm 2.97$ |
| 3 | 80 | $0.29 \pm 0.090$ | $28.05 \pm 1.32$ | $23.79 \pm 1.54$ | $0.80 \pm 0.039$ | $78.00 \pm 5.65$ |
| 4 | 20 | 0.14 ± 0.20 | 28.70 ± 3.32 | 29.67 ± 2.24 | 0.80 ± 0.113 | 69.75 ± 10.05 |
| 5 | 60 | 0.29 ± 0.15 | 28.60 ± 1.58 | 25.37 ± 1.69 | 0.76 ± 0.047 | 77.00 ± 4.13 |
| 6 | 300 | 0.69 ± 0.074 | 27.80 ± 0.72 | 26.57 ± 0.79 | 0.75 ± 0.025 | 70.25 ± 2.86 |
| $\chi^2$ | NA | 148.27 | 0.64 | 71.07 | 21.41 | 98.24 |
| d.f. | NA | 5 | 5 | 5 | 5 | 5 |
| p value | NA | <0.0001 | 0.9861 | <0.0001 | 0.0007 | <0.0001 |

B)
| Year-Season<br>Cluster | n | Density | Temperature | Salinity | Water<br>Depth | SAV |
| --- | --- | --- | --- | --- | --- | --- |
| 1 | 47 | 0.00 ± 0.00 | 29.40 ± 0.25 | 28.12 ± 0.96 | 1.00 ± 0.041 | 53.00 ± 5.94 |
| 2 | 705 | 0.51 ± 0.075 | 25.30 ± 0.48 | 24.55 ± 0.59 | 0.71 ± 0.015 | 70.00 ± 2.10 |
| 3 | 47 | 0.00 ± 0.051 | 29.20 ± 0.28 | 18.26 ± 2.59 | 0.83 ± 0.60 | 59.00 ± 6.51 |
| 4 | 141 | 0.29 ± 0.18 | 30.01 ± 1.20 | 24.58 ± 1.25 | 0.72 ± 0.40 | 66.25 ± 5.36 |
| $\chi^2$ | NA | 89.77 | 32.28 | 70.07 | 85.68 | 22.03 |
| d.f. | NA | 3 | 3 | 3 | 3 | 3 |
| p value | NA | <0.0001 | <0.0001 | <0.0001 | <0.0001 | <0.0001 |

**Fig. 4.**
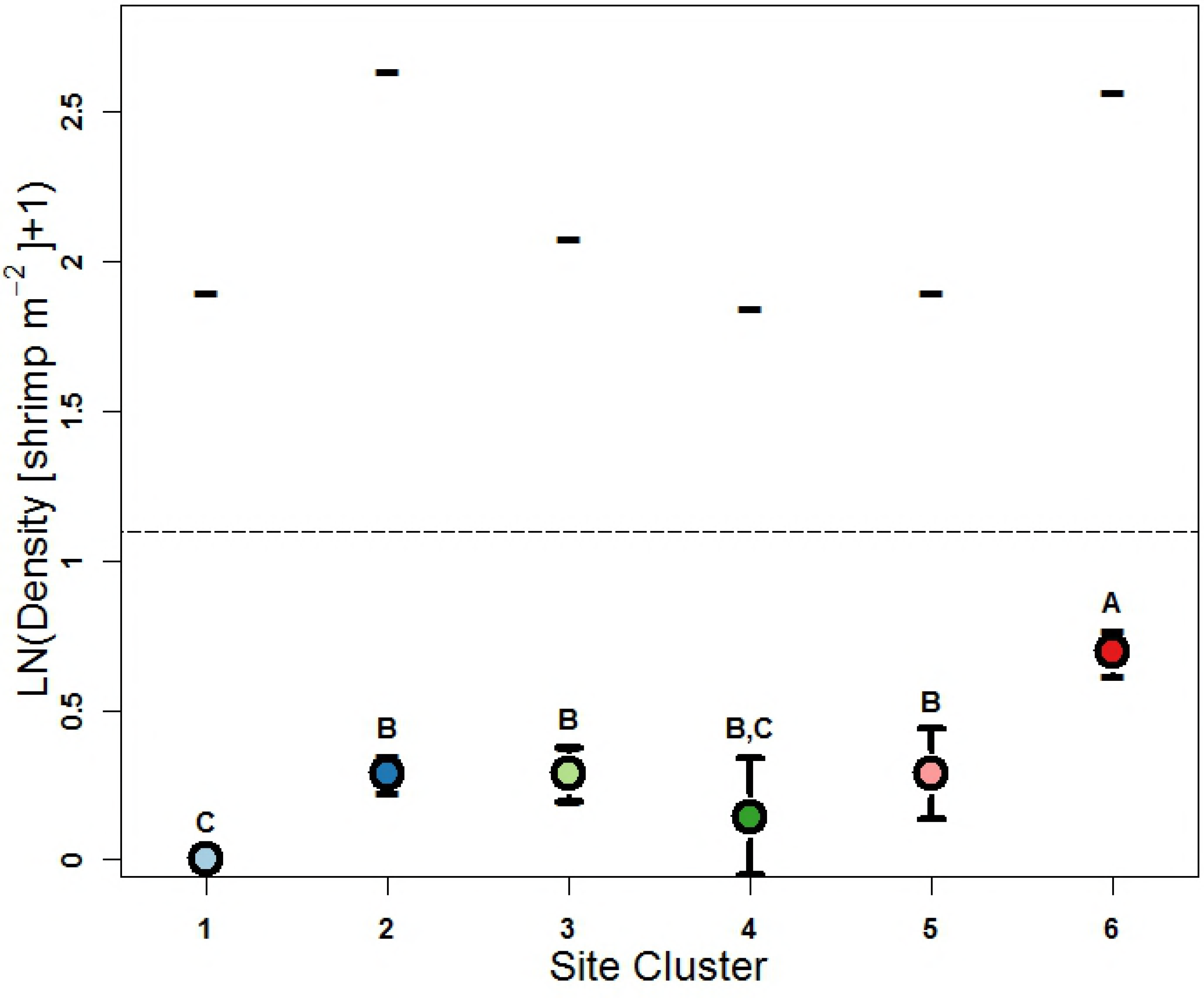
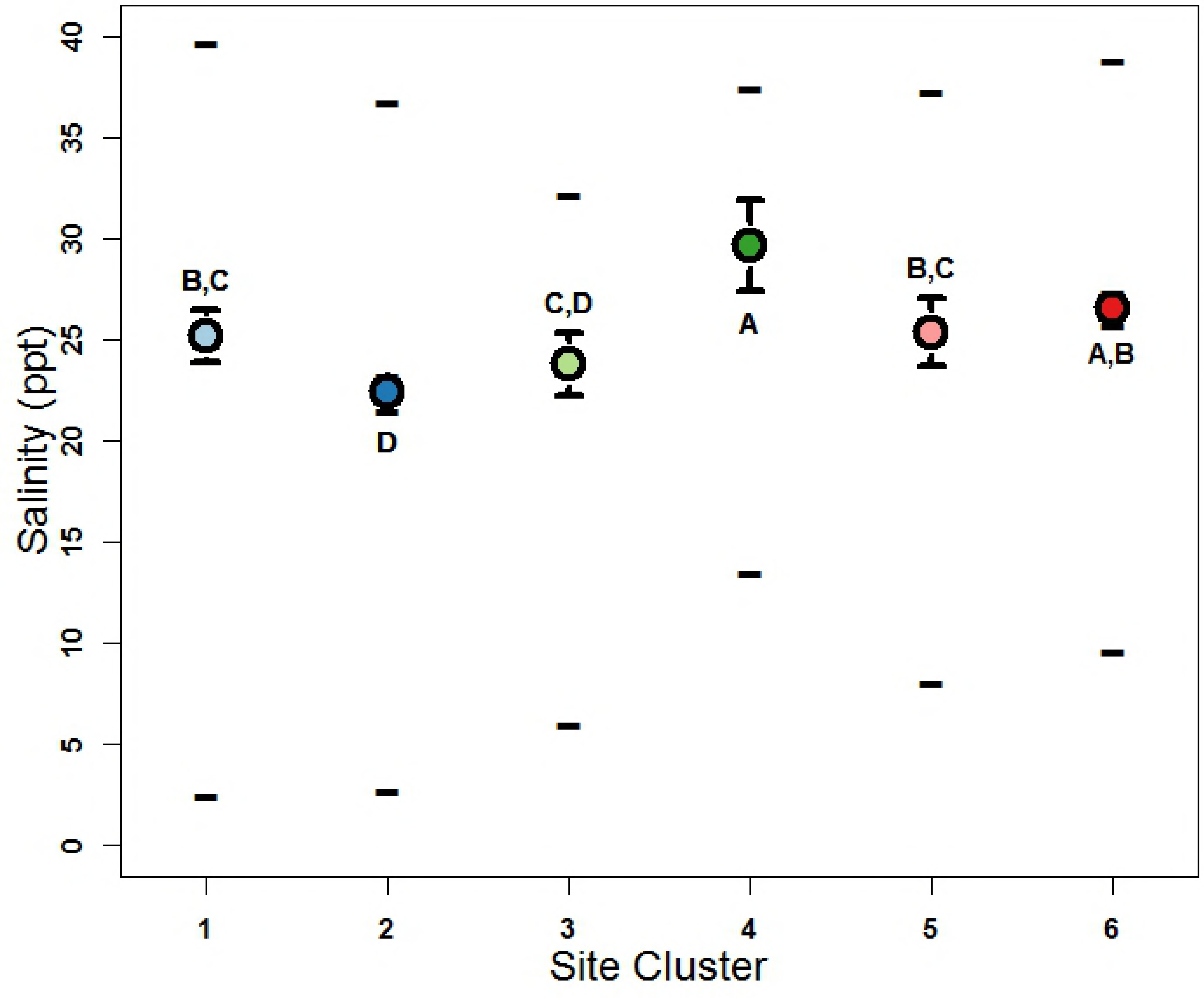
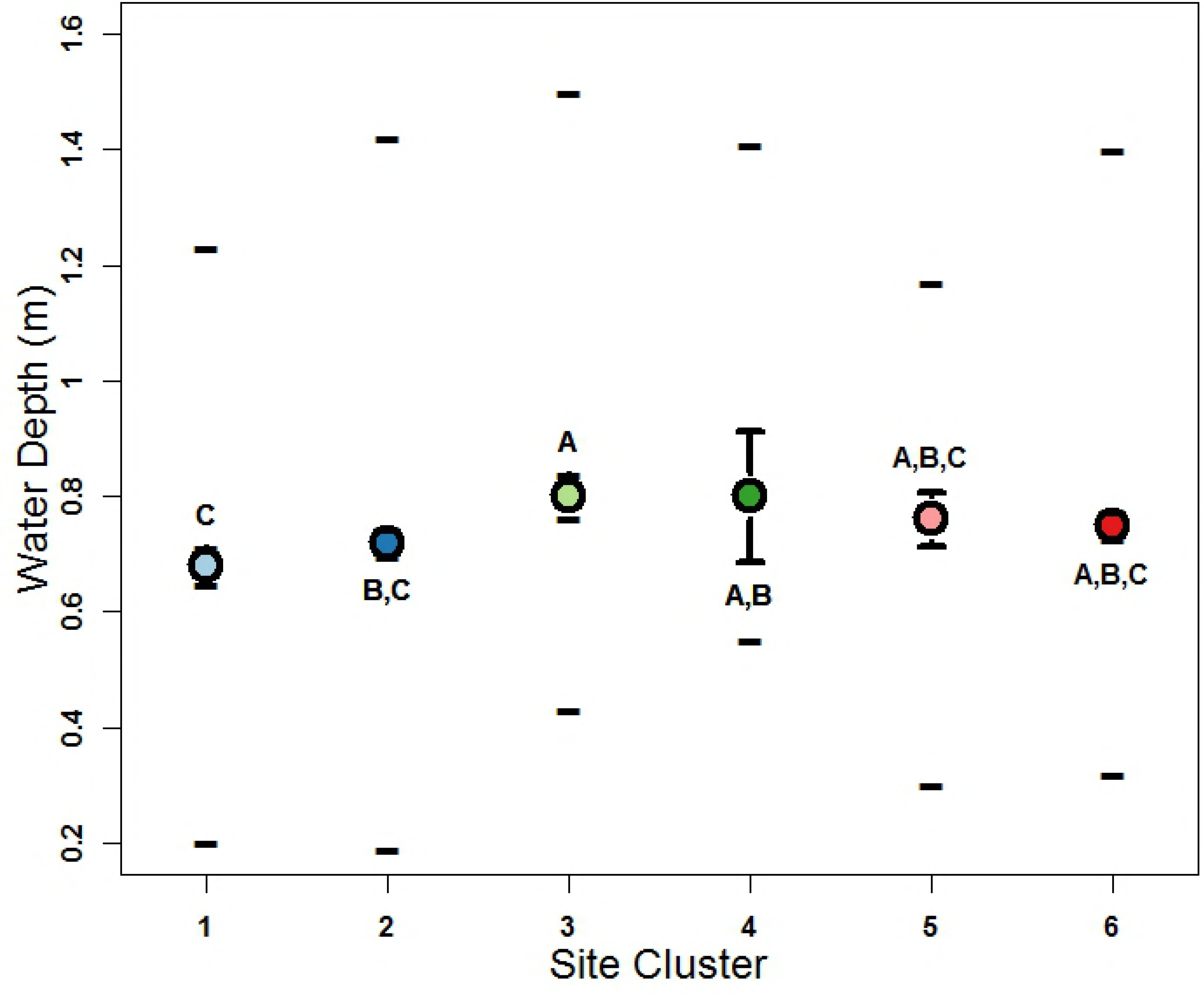
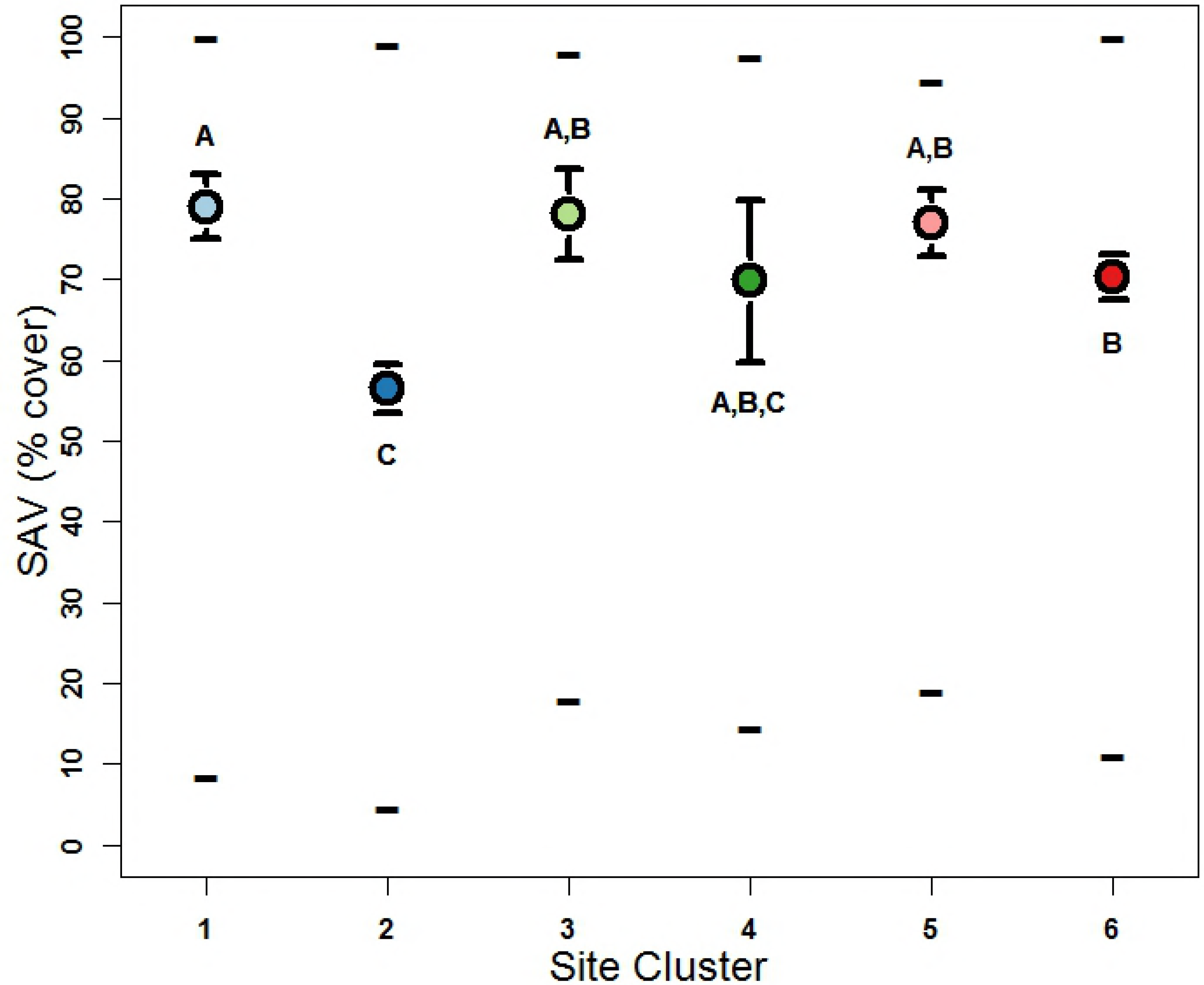
Median (± CI) and maximum, and minimum values of A) density (shrimp m^−2^: LN([x+1]), B) salinity (ppt), C) water depth (m), and D) submerged aquatic vegetation (SAV: % cover) in shrimp density site clusters. Point colors coincide with Fig. 1 and 3B. Letters denote statistically similar groups. Horizontal line of A) depicts the 2 shrimp m^−2^ CERP Interim Goal.

**Fig. 5.**
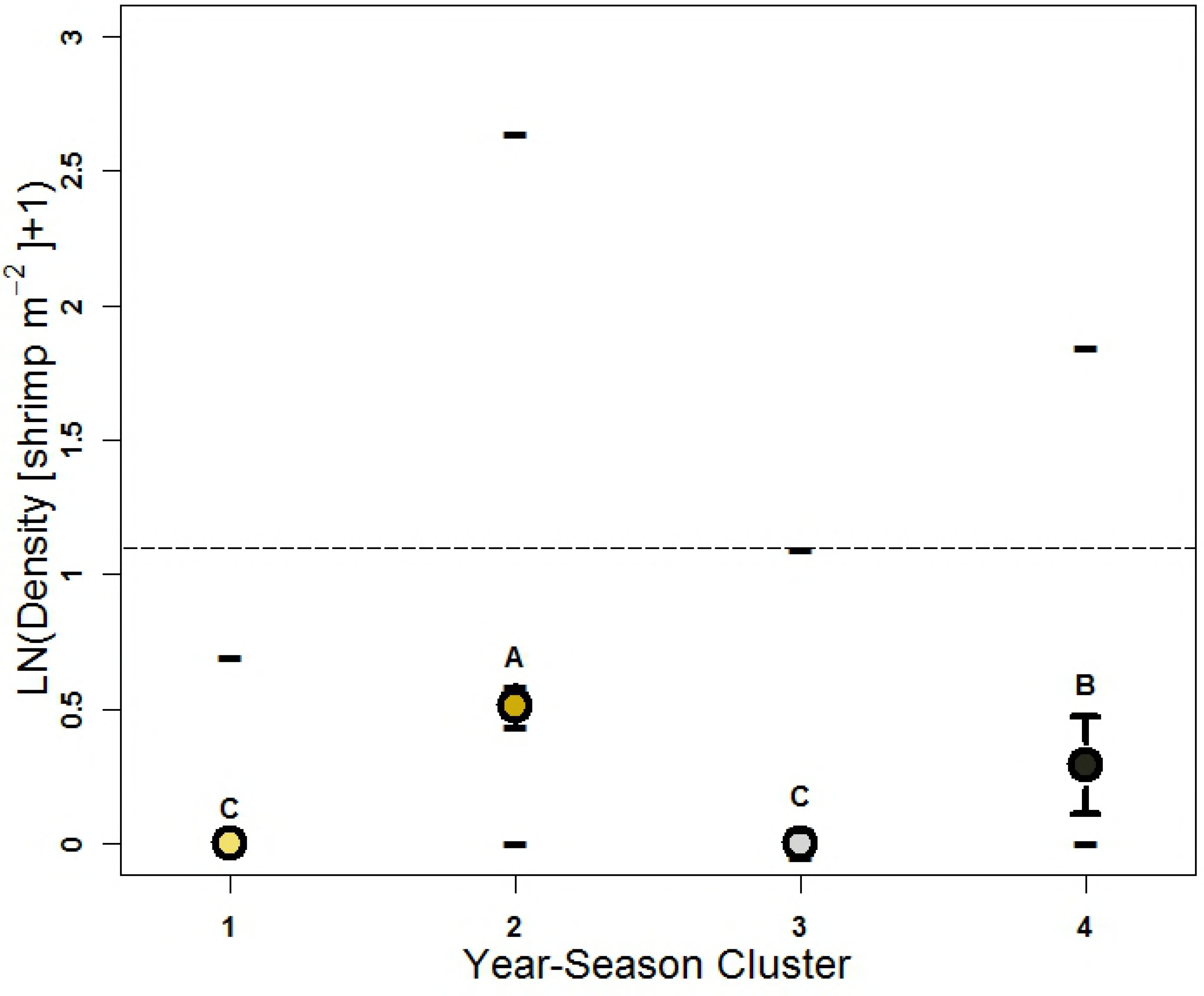
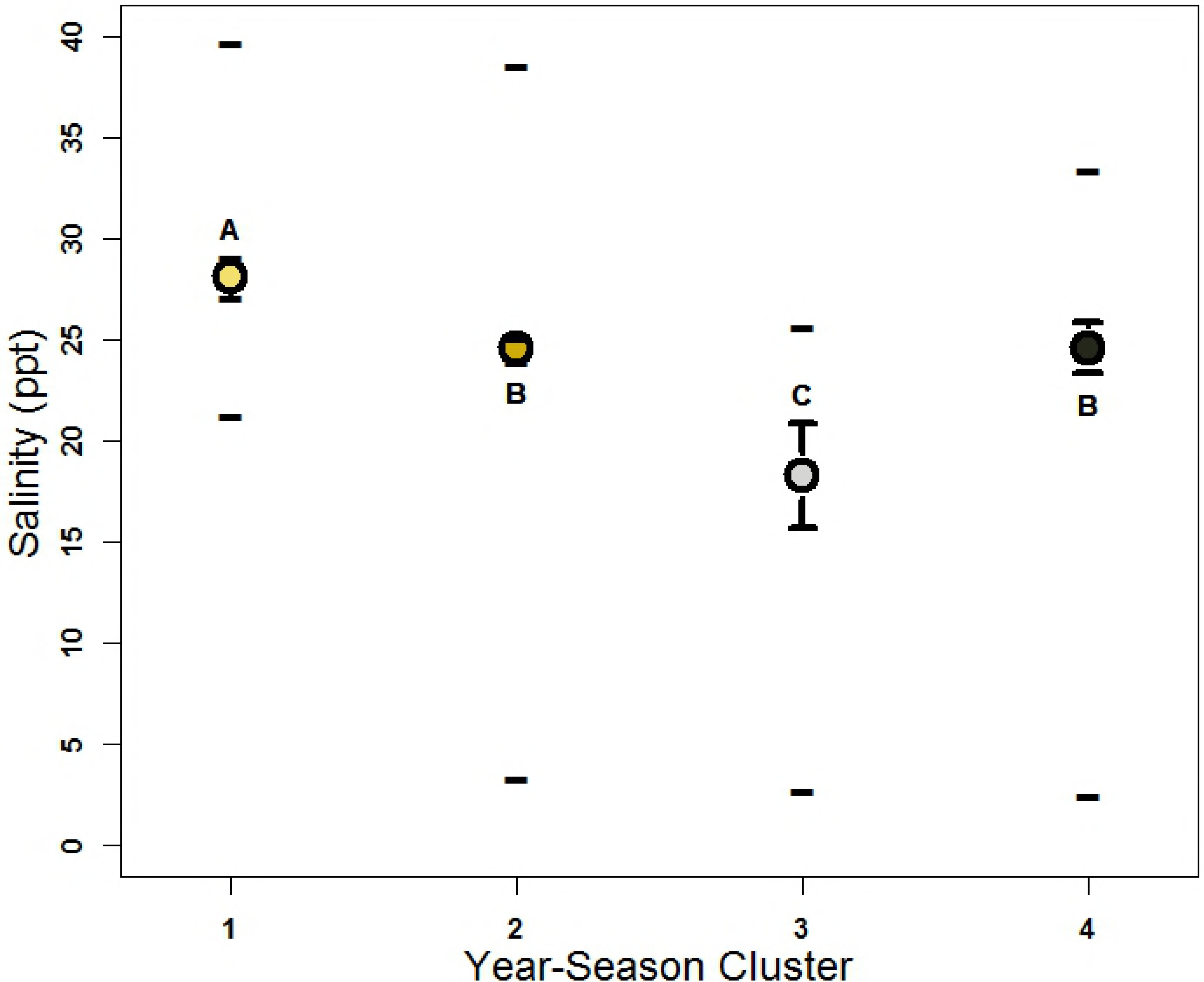
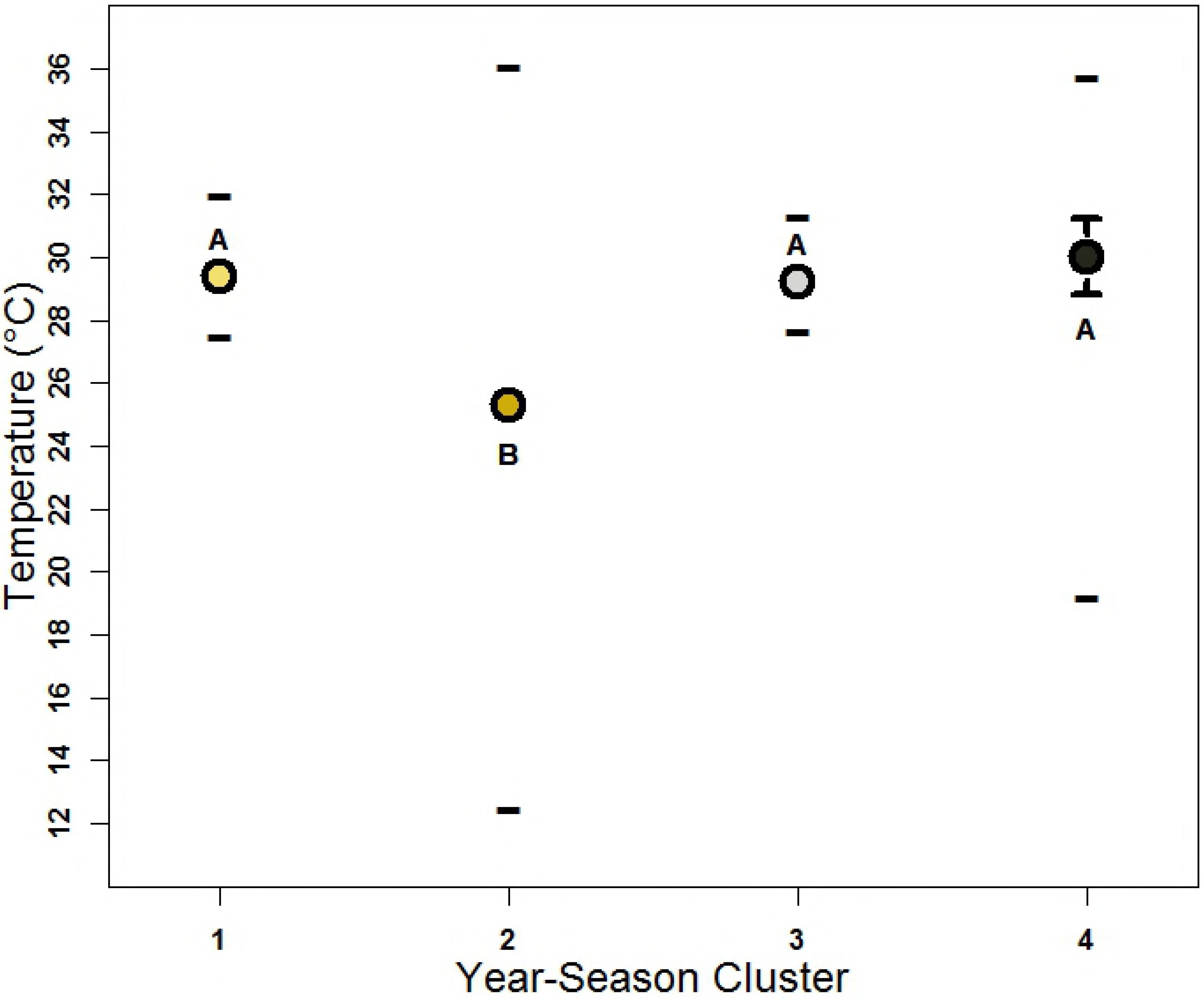
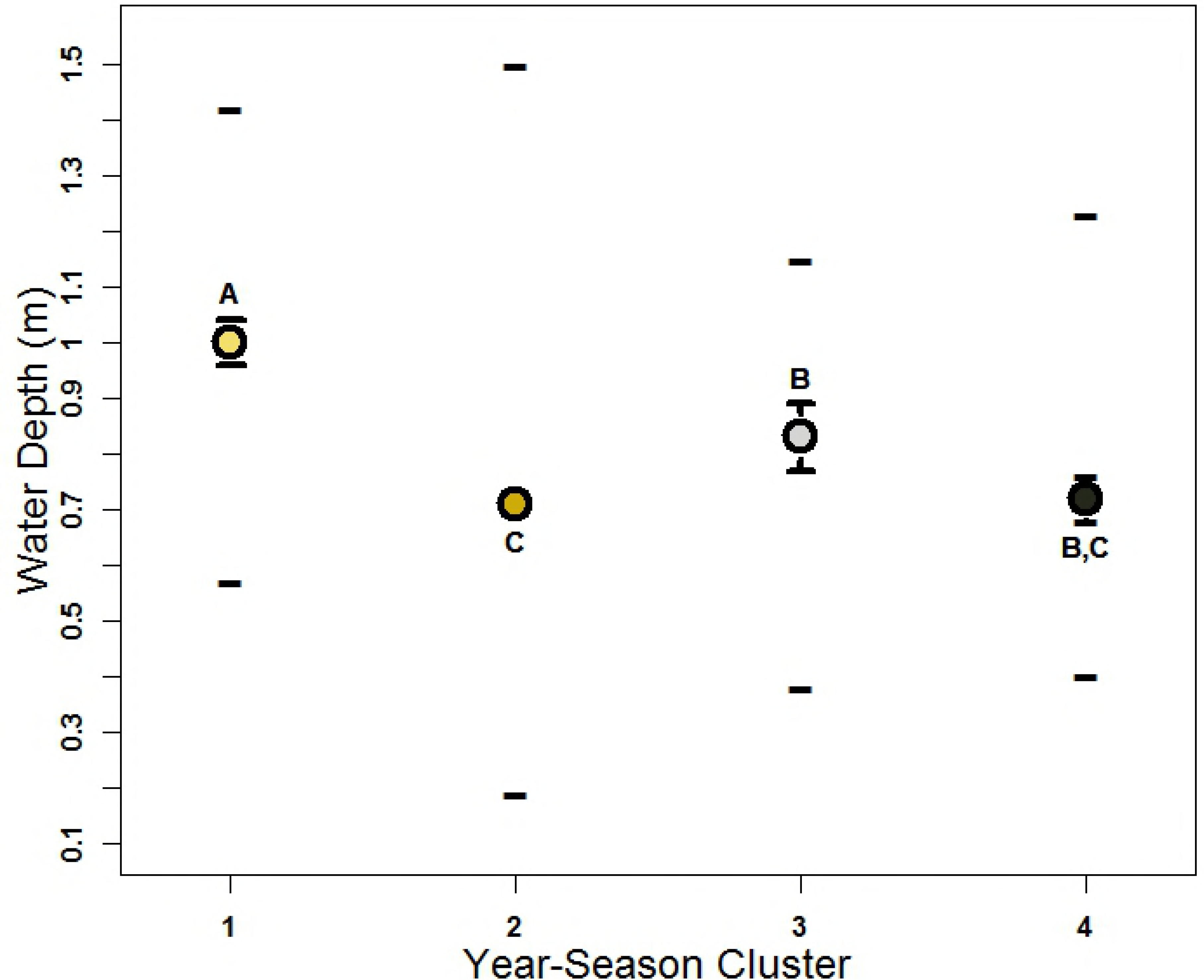
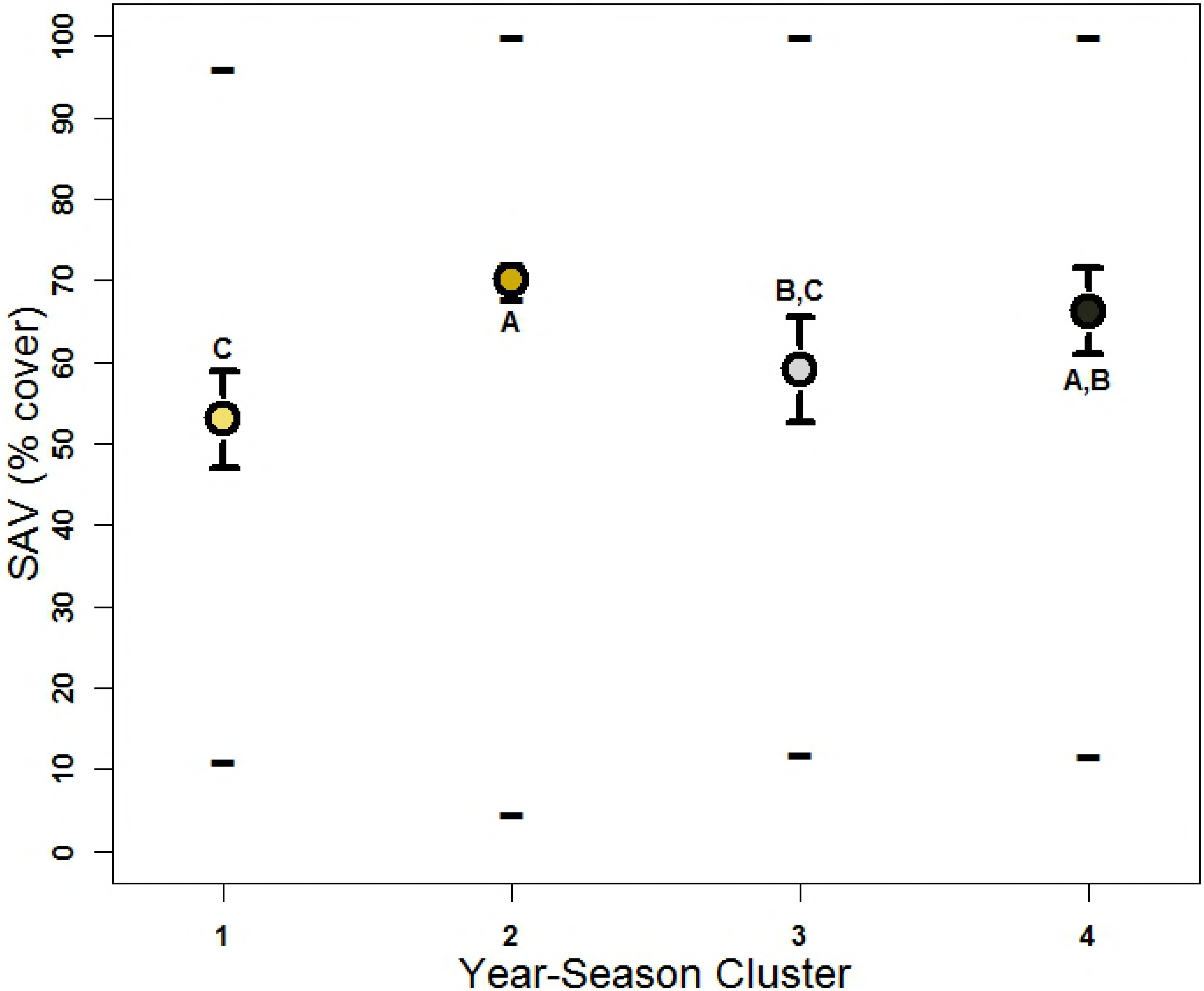
Median (± CI) and maximum, and minimum values of A) density (shrimp m^−2^: LN([x+1]), B) temperature (°C), C) salinity (ppt), D) water depth (m), and E) submerged aquatic vegetation (SAV: % cover) relative to shrimp density year-season clusters. Point colors coincide with Fig. 3B. Letters denote statistically similar groups.

## Discussion

The analysis of 10 years of monitoring data revealed few instances (11.2 %) of pink shrimp densities > 2 shrimp m^−2^, the IG for Biscayne Bay pink shrimp populations [1]. All but one spatially averaged year-season density and all but a few temporally averaged site densities in the 2007-2016 database were significantly below 2 shrimp m^−2^. CERP implementation is expected to result in more favorable salinity conditions for pink shrimp, leading to higher shrimp densities [1,5,27]. Reductions in extreme salinity variability in the southern half of the study area (i.e., Black Point to Convoy Point: Fig. 3.1) could lead to average densities > 2 shrimp m^−2^ across the entire study spatial domain. However, our results suggest that ~ 8ppt salinity (i.e., low mesohaline to oligohaline: Fig. 2B) may have been a threshold below which pink shrimp densities were severely limited. Above this threshold, the log-linear response continued to increase in a more linear fashion. Limitation of pink shrimp densities at salinities <8 ppt does not support coexistence of CERP post-restoration IGs of >2 shrimp m^−2^ and reduction of Biscayne Bay nearshore salinity regimes to oligohaline and low mesohaline conditions.

Spatial pink shrimp density patterns from Black Point to Convoy Point (Fig. 1) were dominated by membership within one low density site cluster (cluster 1) and one intermediate density site cluster (cluster 2). This zone is strongly influenced by canal discharges [11,23–26,33,68]. Both rapid (<60 min to 2 d) and extreme (~25 ppt) salinity reductions can occur along this stretch of coastline [22,24,25,29,69,70]. Such salinity fluctuations can alter fish community assemblages [23] and may affect foraging behavior and survival [22,23]. Pink shrimp may avoid these conditions as they have been reported to migrate to avoid large-volume riverine inflows [71]. Rapid salinity reductions of greater than 20 ppt cause near complete pink shrimp mortality in laboratory settings [72–74]. The low and intermediate density site clusters 1 and 2 included sites 45, 46, and 47 (Fig 3,4), which are located near Turkey Point (Fig 1), well south of the canal zone (Black Point to Convoy Point) and so were not impacted by low and variable salinity conditions. These sites had moderate minimum salinities (≥11.06 ppt), the highest average salinity across all sites (≥29.57 ppt: Table 2), and generally high SAV cover (Table 2, Fig. 3F). The cause of their low shrimp density remains undetermined but does not seem to be related to limitation due to salinity or SAV cover.

Site cluster 6 (Fig. 1), also comprised of sites not located in the canal-zone section of shoreline, had the highest median density. This site cluster had the second highest minimum salinity (9.62 ppt) and was comprised of sites located further from canal mouths (Fig. 1). Previous field [9,77] and modeling [35,76] studies describe the area corresponding to the more northern sites of this cluster as an area of relatively high shrimp abundance. These northern sites were situated immediately across the bay from a large ocean inlet known as the Safety Valve, considered a primary postlarval immigration pathway [35,76], which may have contributed to their high densities [77,78]. Shrimp cumulative size frequency distributions differed between these northern (sites 1 ‑ 17) and southern (sites 18 ‑ 47) sampling sites (Fig S1) with the maximal difference at 7.54 mm CL. This suggests higher abundance of juvenile sized shrimps occurred in the north, which may have been related to growth and/or mortality more so than recruitment. Further, the inclusion of more southern sites (33, 34, 37, 40, 41, 43, and 44) within site cluster 6 (Fig. 1,4) does not fully support the notion of recruitment limitation. These sites are located near mangrove creeks that drain more natural watersheds, although their watersheds are likely reduced due to inland canalization. Several sites, both northern and southern, are located near, but not immediately adjacent to, canals that discharge relatively small volumes of freshwater (Military Canal: 1994-2003 annual mean canal output = 21.9 cfs; Cutler Drain C-100: 1994-2003 annual mean output = 46.1 cfs; [33]). One might anticipate higher shrimp densities along shoreline segments that experience lower-volume freshwater discharges. However, sites immediately adjacent to low-volume discharge canals (i.e., site 6 and site 32) clustered with intermediate and low-density site clusters (cluster 1 and 2, respectively: Fig 3.1, 3.3B, 3.4A).

Most year-seasons (75%) were aggregated within one large cluster, indicating a general lack of inter-annual and inter-season variability in Biscayne Bay juvenile pink shrimp populations. However, the majority (60%) of year-season cluster 2 consisted of dry season sampling events. Observation of higher densities in the dry season is at odds with the pink shrimp IG, which focused on improvement of ‘peak’ fall (wet) season abundances [1]. The small shrimp size associated with this significant difference (5.53 mm CL) suggests that abundance of small juveniles ‑ presumably due to recent postlarval recruitment ‑ caused this seasonal difference. A lack of understanding of Biscayne Bay pink shrimp recruitment complicates study of their abundance patterns. The only study available on recruitment reported a late fall through early winter peak (i.e., October through March: [79]), although the study’s short duration (1 yr) limited consideration of inter-annual trends or consistency. This peak agreed with juvenile abundance studies reporting a late fall/early winter peak [75,80]. Modeling of pink shrimp postlarval recruitment found that oceanographic processes favored Florida Keys potential recruitment during late wet season and early dry season months [81]. Presumably these conditions also favor recruitment to Biscayne Bay: modeling of larval permit *Trachinotus falcatus* originating from spawning grounds near those of pink shrimp also found similar recruitment patterns for the Florida Keys and Biscayne Bay [82]. Oceanographic, coastal, and climatic conditions affect pink shrimp adult reproductive activity [83,84], and larval abundances [85] and interact with behavior to influence early life stage recruitment to nearshore areas [81,86–93]. Use of pink shrimp and other offshore spawning species as indicators of ecological conditions in their nearshore nursery grounds is complicated by life cycles affected by unrelated, prior, external and stochastic conditions [5,32].

Quantile regression found four habitat attributes (water temperature, salinity, water depth, and SAV % cover) that exhibited strong limitation of pink shrimp densities. Two of these (i.e., salinity regime and SAV % cover) can be influenced by freshwater management. Procrustean analysis confirmed the influence of all each habitat attribute except SAV % cover. It should be noted that these habitat attributes vary at differing time scales; for example, water depth can differ by as much as 1.3 m within 12 hr during extreme tidal cycles while SAV % cover may be integrative of salinity, nutrient, water clarity, and other influential factors from 6 mo. to 1 yr or more.

The previously discussed seasonal pattern was confirmed by Procrustean analysis, with QR results indicating higher pink shrimp densities in the dry season and temperature yielding one of the higher concordances of the habitat attributes tested. The pink shrimp IG was based upon observation of a summer/fall (i.e., wet season) peak in abundance [8], which was consistent with Diaz [9], although his investigation was limited to on the four summer/fall months June, July, August, and September. Others reported peak juvenile abundances in late fall/early winter (i.e., November/December) [12] or estimated maximal Biscayne Bay juvenile pink shrimp populations occurring in November (i.e., late fall) [8,75]. Differences in sampling gear and spatial domain and the short durations (≤2 yr) of the four reference studies [8,9,75,80] complicate comparisons with the present study. Although of greater duration, the present study’s bi-seasonal sampling effort may be of insufficient resolution to precisely identify the period of peak pink shrimp density, especially if it changes from year to year.

Procrustean analysis revealed water depth (R^2^ = 28.3) explained the most variability in pink shrimp density of the four habitat attributes presently investigated. This was unexpected given the narrow spatial sampling domain within the mangrove-seagrass ecotone. Associations between nearshore pink shrimp abundance and depth have been previously reported [75,80,94,95]. Other studies that focused on very nearshore areas (<100 m) also found higher abundances there [8–10,12,96]. Recruiting postlarval pink shrimp often concentrate in SAV near the low-tide mark along shorelines [9,12,97–102]. Fluctuating tidal depths and/or detection probability at greater depths likely contributed to the domed shape of the QR relationship [103]. Procrustean analysis confirmed a direct relationship between salinity and density, but also suggested that salinity was less influential than temperature (i.e., seasonality) or water depth on spatiotemporal density patterns. Other pink shrimp habitat investigations [80,94] found multiple habitat attributes (e.g., salinity, salinity standard deviation, standard deviation of turbidity, temperature, median sediment size, dissolved oxygen concentration, water depth, and benthic habitat characteristics) can influence pink shrimp abundance. As re-iterated by Zink et al. [6], Costello et al. [12] stated that “….factors other than salinity per se control abundance of the euryhaline juveniles...”

Re-analyses by Zink et al. [6] of data presented by Brusher and Ogren [104] and Minello [105] found increasing abundance with increasing salinity and no statistical difference between polyhaline and mesohaline pink shrimp abundances. It was unexpected to not find a limitation of pink shrimp density at higher salinities, especially at conditions >35 ppt (e.g., a significant, negative quadratic term or the significant cubic splines functions in Fig. A2, Table A1). The range of salinity values observed in the study did not include hypersaline values (>40 ppt). Perhaps this range of salinity observations was not broad enough to characterize suspected reduction of densities in extreme hypersaline conditions [5].

The Biscayne Bay pink shrimp IG suggested a pink shrimp preference for seagrasses, and presumed that increased % cover of seagrasses would increase pink shrimp abundance [1,5,30] and increase the seaward spatial extent of *H. wrightii* [30]. Presently, total SAV QRs yielded the most plausible relationship between pink shrimp density and the benthic habitat metrics investigated. Contrary to this relationship, associated Procrustean analysis test results were non-significant. The seemingly weak statistical relationships with either total or species-specific SAV metrics was unexpected. Pink shrimp associations with *H. wrightii* have been previously reported [12,96,97,106], while other studies have reported maximal pink shrimp densities relative to total SAV biomass or % cover [10,107,108]. Although one study reports negative impacts of drift and attached algal biomass [109], the positive relationships reported by most studies suggested a stronger relationship between pink shrimp density and either species-specific or total SAV would be readily observed.

Several environmental perturbations occurred during this study. Variability in climatic conditions led to both wetter and drier than normal wet seasons (Fig. 3D). However, alteration of typical salinity regimes did not seem to influence temporal density patterns substantially. For example, the second highest wet season pink shrimp density (1.29 ± 1.65 shrimp m^−2^: Table 1) coincided with 2012 record rainfall that reduced salinities (3.34 to 22.08 ppt) across the spatial domain (Fig. 3D). Conversely, the highest wet season pink shrimp average density (1.45 ± 2.25 shrimp m^−2^: Table 1) occurred during the 2015 wet season, a period that was previously denoted as a ‘hypersaline’ period [65]. Record dry season rainfall during 2016 yielded the lowest average dry season salinity (Table 1, Fig. 3D) while pink shrimp densities that dry season were moderate (0.84 ± 1.18 shrimp m^−2^: Table 1). Despite their differing salinity conditions, these year-seasons were assigned to the same shrimp density cluster (Fig. 3B). The 2013 wet season clustered separately from the others, suggesting a negative impact of microalgal bloom conditions [65,66] on pink shrimp density. No discernable impact on pink shrimp densities was observed due to passage of an extreme cold front in the 2010 dry season.

Due the field sampling design, the present results apply only to shallow nearshore areas very near (mainly within ~50 meters of) the shoreline. Application of the present results to areas further offshore should proceed with caution, if done at all, due to potential interaction with other habitat attributes that influence trends in pink shrimp density. The study was also limited by apparent low catchability of very recently settled pink shrimp by the throw trap gear manifested as reduced numbers of pink shrimp from 3 to 5 mm CL (Fig. S1). Pink shrimp postlarvae are generally considered settled in their nursery habitat by 3 mm CL [9,12,86–88]. Pink shrimp postlarvae settle in the shallow (≤1 m), calm water areas along shorelines [12,106], which would suggest they should be readily available to the present field sampling program that samples nearshore waters generally < 1 m deep.

The RECOVER Biscayne Bay IG anticipates >2 shrimp m^−2^ as a target wet season pink shrimp density to be achieved with CERP BBCW implementation. But achievement of BBCW and CERP salinity IGs, which include low mesohaline (<10 ppt) and even oligohaline conditions (<5 ppt), may negatively impact pink shrimp density. The Biscayne Bay pink shrimp IG may need modification to clarify whether the ≥ 2 shrimp m^−2^ target refers to all monitoring observations or a seasonal or annual average density across the entire shoreline and to further consider spatial and seasonal abundance patterns.

## Acknowledgements

This study used monitoring data collected over a substantial period by many of individuals; without their efforts, this study would not have been possible. This study was conducted during pursuit of the PhD degree of I. Zink; he would like to thank committee members D. Die and J. Luo for their comments and suggestions during review of earlier versions of this manuscript. The authors would also like to thank two anonymous reviewers for their suggestion prior to submission of the manuscript for publication.

